# Modern spandrels: The roles of genetic drift, gene flow and natural selection in the formation of parallel clines

**DOI:** 10.1101/289777

**Authors:** James S. Santangelo, Marc T. J. Johnson, Rob W. Ness

**Author notes:** Author for correspondence: James S. Santangelo. Electronic supplementary material is available online at https://dx.doi.org/XXX.

## Abstract

Urban environments offer the opportunity to study the role of adaptive and non-adaptive evolutionary processes on an unprecedented scale. While the presence of parallel clines in heritable phenotypic traits is often considered strong evidence for the role of natural selection, non-adaptive evolutionary processes can also generate clines, and this may be more likely when traits have a non-additive genetic basis due to epistasis. In this paper, we use spatially-explicit simulations modelled according to the cyanogenesis (HCN) polymorphism in white clover (*Trifolium repens*) to examine the formation of phenotypic clines along urbanization gradients under varying levels of drift, gene flow and selection. HCN results from an epistatic interaction between two Mendelian-inherited loci. Our results demonstrate that the genetic architecture of this trait makes natural populations susceptible to decreases in HCN frequencies via drift. Gradients in the strength of drift across a landscape resulted in phenotypic clines with lower frequencies of HCN in strongly drifting populations, giving the misleading appearance of deterministic adaptive changes in the phenotype. Studies of heritable phenotypic change in urban populations should generate null models of phenotypic evolution based on the genetic architecture underlying focal traits prior to invoking selection’s role in generating adaptive differentiation.

## Introduction

Evolutionary clines—changes in the frequency of a genotype or a heritable phenotype over a geographical area [1]—have long served as model systems in evolutionary biology [2]. Clines arise and are maintained via the interplay of genetic drift, gene flow and natural selection across an environmental gradient [3–5]. The multiple evolutionary mechanisms structuring clines has prompted their continued use by evolutionary biologists seeking to explore the relative contributions of non-adaptive and adaptive evolutionary processes in structuring patterns of genetic and phenotypic diversity and differentiation within and between populations.

While clines are often interpreted as strong evidence of adaptive evolution, non-adaptive processes (e.g. genetic drift and gene flow) may also generate correlations between phenotype frequencies and environmental gradients. For example, local genetic drift in combination with spatially restricted gene flow (i.e. isolation by distance, [6]) can generate clines in allele frequencies at a single locus [7]. Similarly, serial founder events can generate clines in quantitative traits controlled by multiple loci [8] and phenotypic clines can arise through multiple introductions from a species’ native range during invasion [9]. Disentangling the relative importance of stochastic and deterministic forces in the formation of clines is thus essential prior to invoking selection’s role in generating differentiation among populations.

If multiple independent clines in the same direction. (i.e. parallel clines) at single-loci or in quantitative traits arise by neutral processes, then clines in either direction should occur in direct proportion to the initial frequency of underlying alleles. For example, if the initial frequency of underlying alleles is 0.5, then each locus is equally likely to drift either upward or downward, resulting in no consistent change in allele frequency across clines. Thus, the presence of parallel clines is strong evidence for the role of natural selection, as putative adaptations are unlikely to evolve repeatedly in the same direction via stochastic forces [10]. However, when traits have a non-additive genetic basis due to epistasis, clines may occur more frequently in a particular direction because stochastic changes in allele frequencies at one locus can have a disproportionate effect on the distribution of phenotypes within a population. For example, stochastic forces have caused the repeated loss of the Mendelian inherited, epistatically determined short-style (S) morph from tristylous populations of *Eichhornia paniculata* in northeastern Brazil, Jamaica, and Cuba [11–13]. The fact that drift can lead to directional changes in non-additive traits across multiple, independent populations means that the presence of parallel clines in such traits is insufficient evidence for the role of selection in generating adaptive differentiation. In such cases, selection should only be invoked upon observing either a greater frequency of clines, or stronger clines, than would be expected by the effects of drift alone. Studies exploring genetic and phenotypic evolution across replicated environmental gradients, while considering the genetic architecture of the trait in question, would provide a strong test of the relative contributions of drift, gene flow and selection in the formation of parallel clines.

Urbanization is one of the most widespread human disturbances on earth and it provides an excellent large-scale replicated system to understand how adaptive and non-adaptive evolutionary processes contribute to the formation of parallel clines. Changes in biotic and abiotic factors associated with urbanization has led to parallel evolutionary responses in taxa as diverse as plants [14], fish [15], lizards [16], birds [17], insects [18], and humans [19]. In addition, the widespread fragmentation associated with urbanization has resulted in gradients in the strength of genetic drift and gene flow. For example, urban fragmentation has reduced genetic diversity and increased differentiation among urban populations of white-footed mice (*Peromyscus leucopus*) in New York City [20, 21], and eastern red-backed salamanders (*Plethodon cinereus*) in Montréal, Canada [22], due to increased genetic drift and reduced gene flow among urban populations. Despite these advances, the relative roles of genetic drift, gene flow and selection in affecting the evolution of populations in urban environments is presently unknown.

Recently, Thompson *et al.* (2016) identified parallel urban-rural clines in the frequency of plants producing hydrogen cyanide (i.e. cyanogenesis, HCN)—a potent antiherbivore defence— in populations of white clover (*Trifolium repens*) across multiple cities. They found that HCN defended genotypes were less frequent in urban populations in 3 of the 4 cities examined [23]. The authors identified lower winter surface temperatures in urban populations as a putative selective agent structuring urban-rural cyanogenesis clines. However, they did not investigate the alternative hypothesis that these clines could be caused by genetic drift, which is especially likely given the epistatic genetic architecture underlying cyanogenesis (figure 1). In this study, we use the HCN polymorphism in white clover as a model for exploring the conditions under which non-adaptive (i.e. genetic drift and gene flow) and adaptive (i.e. selection) processes can generate repeated clines in phenotypes with an epistatic genetic basis. We address the following specific questions: (1) Can genetic drift influence the formation of spatial clines in HCN? (2) How does selection affect the occurrence and strength of spatial clines in HCN? (3) What are the interactive effects of genetic drift and selection in the formation of clines in HCN? In all simulations used to address the questions above, we manipulated levels of dispersal to examine the homogenizing effects of gene flow on cline formation. We discuss the role of adaptive and non-adaptive evolutionary processes in the evolution of cyanogenesis clines in white clover along urbanization gradients and other environmental gradients and more broadly with respect to the role of genetic drift in the formation of parallel clines.

**Figure 1:**
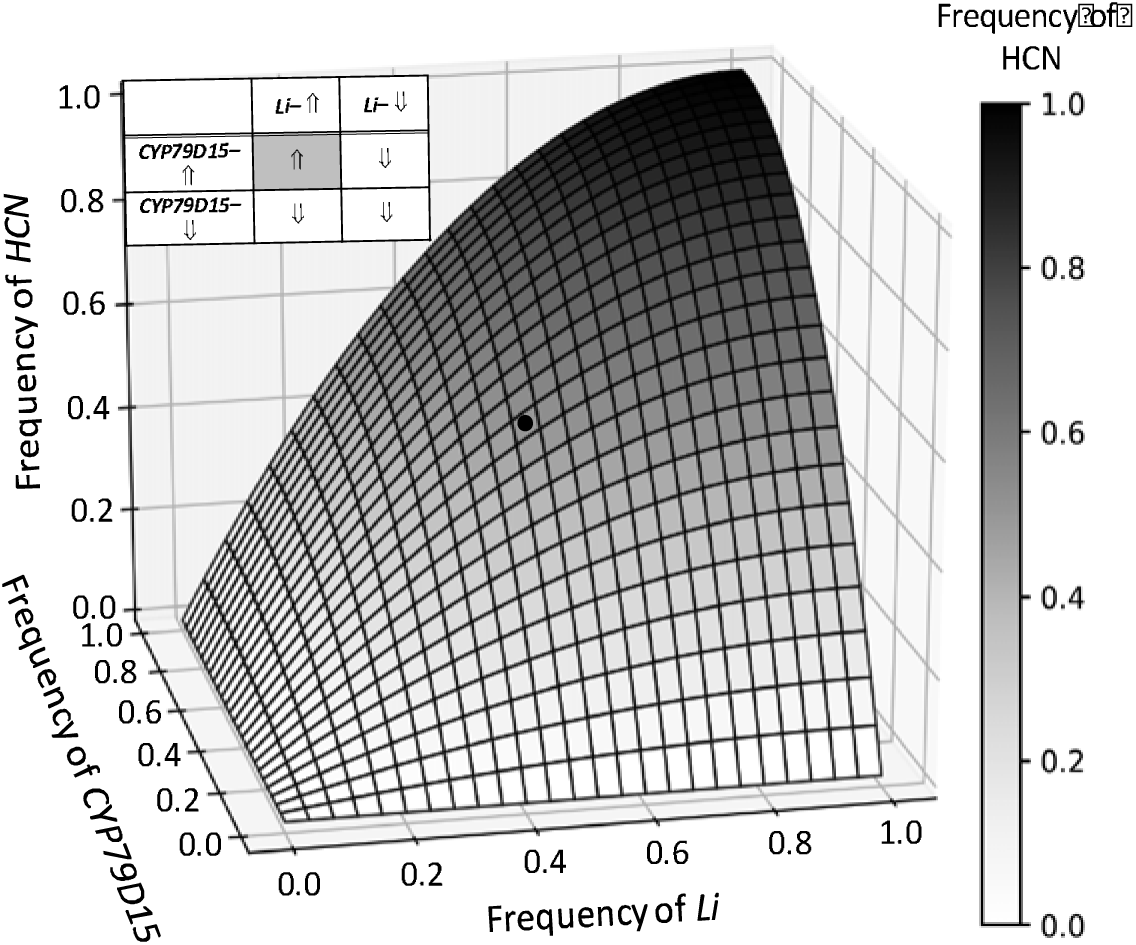
Changes in the frequency of the cyanogenic phenotype (HCN) with changes in the frequency of either dominant allele (i.e. *CYP79D15* or *Li*). HCN is controlled by two independently segregating (i.e. unlinked) Mendelian loci (*CYP79D15* and *Li*) and plants require a dominant allele at both loci to produce HCN. The duplicate recessive epistasis (i.e. complementary epistasis [49]) underlying the production of HCN has consequences for how drift is expected to influence the frequency of HCN in natural populations. The black dot represents the point at which the frequencies of both dominant alleles are at 0.5. When both loci are drifting, the frequency of HCN will only increase if the frequency of both underlying dominant alleles increases (table inset). By contrast, a decrease in the frequency of the dominant allele at either locus typically results in a rapid decrease in the frequency of HCN. Thus, the loss of dominant alleles in this system has a greater effect at reducing HCN frequencies than gains in dominant alleles have at increasing HCN frequencies. This makes the loss of HCN from populations more likely solely due to drift.

## Methods

### Overview of simulations

To examine the formation of spatial clines in HCN, we created a series of spatially-explicit simulations in Python (v. 2.7) to track the frequency of alleles at loci underlying cyanogenesis (i.e. *CYP79D15* and *Li,* figure 1) within populations through time and across space. The frequency of HCN within each population is easily calculated from the frequency of recessive alleles at underlying loci as: 1 – [q_CYP_ + q_Li_ - (q_CYP_ × q_Li_)], where q_CYP_ and q_Li_ represent the frequency of the recessive alleles at *CYP79D15* and *Li*, respectively. We represented a transect from urban to rural habitats as a one-dimensional, linear matrix with 40 cells, consistent with the number of populations sampled across cities by Thompson *et al.* (2016). Each cell (hereafter patch) represents a patch of suitable habitat that can support a population of *Trifolium repens*. These simulations allowed for fine scale, independent control of both stochastic and deterministic parameters important for varying and maintaining the frequency of *CYP79D15* and *Li*—and thus HCN—in patches distributed across the landscape (table S1). The order of events in the simulations were as follows: (1) Local reproduction and population growth, (2) selection, (3) gene flow, and (4) colonization.

We simulated a total of eight biologically informed scenarios (table S2): six that explore the effects of drift, gene flow, and selection on the formation of phenotypic and genetic clines in cyanogenesis, and two that examine the effects of drift and initial allele frequencies on clines. We started by assessing the effects of genetic drift on the formation of cyanogenesis clines in the absence of selection. We manipulated drift in two ways: (1) by imposing a gradient in the carrying capacity of populations across the landscape, and (2) through serial founder events. Both scenarios were simulated independently and produced qualitatively similar results (figure S1). We thus focus on drift scenario 1 (i.e. gradient in carrying capacity) and consider drift scenario 2 in the online supplementary materials (methods: text S1; results: text S2, figures S1 to S5, figure S7). In addition, we focus our results on cases where the initial frequency of dominant alleles was 0.5, which resulted in the strongest clines, and consider the effects of initial allele frequency in the electronic supplementary material (text S3, figure S6 and S7).

### Question 1: Does genetic drift influence the formation of spatial clines in HCN?

The first scenario (i.e. gradient in carrying capacity) represents a case where clover populations were initially similar but increased fragmentation associated with urbanization reduced urban population sizes and increased the strength of drift. We imposed a gradient in the carrying capacity (*K*) of populations across the landscape, thereby placing an upper-limit on the population size (*N*, figure S8*a*). Drift is expected to be strongest in smaller populations and this method has been used in other simulations exploring the effects of drift, gene flow, and selection on patterns of local adaptation [24]. We first simulated a scenario where *N* is assumed to be greatest in rural populations (maximum *N* = 1000) and, for simplicity, declines linearly with increasing urbanization (figure S8*a*). We simulated multiple minimum urban population sizes (minimum *N =* 10; 100; 500; 1000), which represented strong to no gradients in drift, respectively. All populations were started at their individual *K*, and they were kept at this size throughout the simulations. These simulations were run for 500 non-overlapping generations. We did not include gene flow in these initial simulations as we sought to explore the formation of clines due to drift alone, independent of other evolutionary mechanisms.

To examine the effects of gene flow on the formation of spatial clines in HCN formed by genetic drift, we performed additional simulations under varying levels of gene flow. To provide the greatest sense of the effects of gene flow, we restricted our simulations to the case of a strong gradient in drift (i.e. rural *N* = 1000, urban *N* = 10), which we expected to generate the strongest clines in HCN. We simulated 13 levels of gene flow (*m* = 0; 0.001; 0.0025; 0.005; 0.01; 0.02; 0.035; 0.05; 0.1; 0.2; 0.35; 0.5, 1.0, table S2) to explore how a wide range in levels of gene flow affects the formation and maintenance of clines via drift. While each population can receive migrants from all populations in a given generation, these values represent the maximum proportion of alleles exchanged between any two populations, which occurs among adjacent populations (text S4). In subsequent simulations, we simulated three levels of gene flow: *m* = 0, 0.01, and 0.05, representing no, low, and high gene flow, respectively, and corresponding to levels of gene flow that resulted in substantial decreases in the strength of clines in the drift scenario described here. Specific details on how we modelled gene flow are described in the online supplementary materials (text S4).

### Question 2: How does selection affect the occurrence and strength of spatial clines in HCN?

We used two-locus selection models to explore the effects of selection in generating and maintaining cyanogenesis clines [25]. Selection favoured either cyanogenic (i.e. HCN+) or acyanogenic (i.e. HCN–) genotypes, depending on the population’s position in the landscape. In our model, selection favouring HCN+ genotypes in rural environments changed gradually along an urbanization gradient until HCN– genotypes were favoured in the urban core, consistent with the phenotypic clines reported by Thompson *et al.* ([23]) and Johnson et al (this issue, [26]). For each simulation, we defined a maximum strength of selection that favoured HCN+ genotypes in the rural-most population and HCN– genotypes in the urban-most population. The selection coefficient varied linearly across the matrix such that HCN+ and HCN– genotypes had equal fitness in the central population of the landscape (i.e. population 20, figure S9). We simulated 10 different maximum selection coefficients (*s* = 0; 0.001; 0.0025; 0.005; 0.0075; 0.01; 0.025; 0.05; 0.1; 0.2) to examine the effects of fine-scale variation in the strength of selection on the formation of clines. For example, a selection coefficient of 0.05 corresponded to a 5% reduction in fitness of HCN+ genotypes in the urban-most population and a 5% increase in fitness of HCN+ genotypes in the rural-most population. These simulations did not include gradients in the strength of genetic drift; rather, all populations in the landscape started—and remained—at *N* = 1000. However, for each selection coefficient above, we simulated three levels of gene flow: *m* = 0, 0.01, and 0.05, representing no, low, and high gene flow, respectively. Thus, these simulations explored the combined effects of selection and gene flow on the formation of clines in HCN in the absence of drift.

When selection acts on two or more loci, linkage disequilibrium (LD) may evolve as genotypes with particular allele combinations are favored, resulting in gamete frequencies that differ from their expectation based on allele frequencies [25]. However, given that the *CYP79D15* and *Li* loci are physically unlinked [27], theory predicts that free recombination (recombination fraction = 0.5) between these loci would limit the accumulation of significant LD, even under selection [28]. Simulations exploring the build-up of LD under varying selection regimes acting for or against cyanogenic genotypes confirmed that even strong selection (*s* = 0.1) results in little accumulation of LD between loci underlying cyanogenesis (text S5, figure S10). We therefore ignored the effects of LD in our simulations and gamete frequencies each generation were calculated directly from allele frequencies, with recombinant gametes being produced with equal frequency (0.25) from heterozygous genotypes.

### Question 3: What are the interactive effects of genetic drift and selection on the formation of clines in HCN?

We sought to understand the combined effects of drift, gene flow and selection on the formation of clines in HCN, and specifically the extent to which selection can counter the formation of clines under drift. We first imposed a gradient in the strength of drift as described above (see “Question 1”), but with the gradient running in the opposite direction: the urban-most population was maintained at a size of *N* = 1000 while the rural-most population was maintained at *N =* 10. Again, we focus on drift scenario 1 because the results from drift scenario 2 were qualitatively similar and are presented in the online supplementary materials (methods: text S1*b*; results: text S2*b*, figure S5). Selection favoured HCN+ genotypes in rural populations and HCN– genotypes in urban populations, as described above. As such, the stochastic loss of dominant alleles in smaller rural populations is countered by their higher fitness. We additionally included three levels of gene flow: *m* = 0, 0.01, and 0.05, representing no, low, and high gene flow, respectively. These simulations enabled us to identify the strength of selection necessary to counteract the loss of HCN due to drift and examine the formation of HCN clines under opposing forces of drift and selection, with varying levels of gene flow.

### Analyses

We performed 1000 iterations for each of the simulated scenarios listed in table S2. For each iteration, we ran a linear regression using within-population HCN frequency as the response variable and distance from the urban-most population (i.e. patch 40) as the predictor. For simulations involving complete colonization of the landscape (i.e. drift scenario 1 above), this regression was performed using HCN frequencies at generation 250, consistent with the age of many large north American cities. Note however that our results are not contingent on the generation chosen for analysis as any generation following the initial formation of a cline until generation 500 produced qualitatively similar results (text S6, figure S11). For simulations involving serial founder events (i.e. drift scenario 2), we ran this regression in the first generation after the entire landscape became filled with populations, though as before running the regression in other generations yielded similar results. In both cases, each regression can have one of three possible outcomes: (1) A positive cline, representing significantly (*P* < 0.05) higher rural than urban HCN frequencies. These clines are consistent in direction with the urban-rural cyanogenesis clines reported by Thompson et al. (2016) and Johnson et al. (2018, this issue); (2) a negative cline, representing significantly higher urban than rural HCN frequencies, and (3) no cline (i.e. *P* > 0.05). For each simulated scenario, we report the proportion of significantly positive and negative clines in addition to the mean slope (β) across all 1000 iterations, independent of significance. To answer our questions, we explore how these proportions and the mean slope are affected by varying levels of genetic drift, gene flow and selection. All analyses were performed in R version 3.4.3 [29].

## Results

### Question 1: Does genetic drift influence the formation of spatial clines in HCN?

In the absence of selection and gene flow, a gradient in carrying capacity from rural (large *N*) to urban (small *N*) disproportionately resulted in the formation of positive clines (i.e. less HCN in urban populations). The mean slope of clines across all 1000 simulations was always positive in the presence of spatial gradients in drift, reaching a maximum of β ≈ 0.007 in the presence of a strong gradient in drift (urban *N* = 10) and becoming gradually weaker as the strength of the gradient was reduced (figure 2*a*). Similarly, ∼28% of clines were significantly positive under strong drift gradients, whereas significantly negative clines (i.e. more HCN in urban populations) were completely absent; significantly negative clines only occurred in the absence of a gradient in drift (urban *N* = rural *N* = 1000), in which drift resulted in an equal frequency of positive and negative clines at a frequency of ∼3% (figure 2*c*).

**Figure 2:**
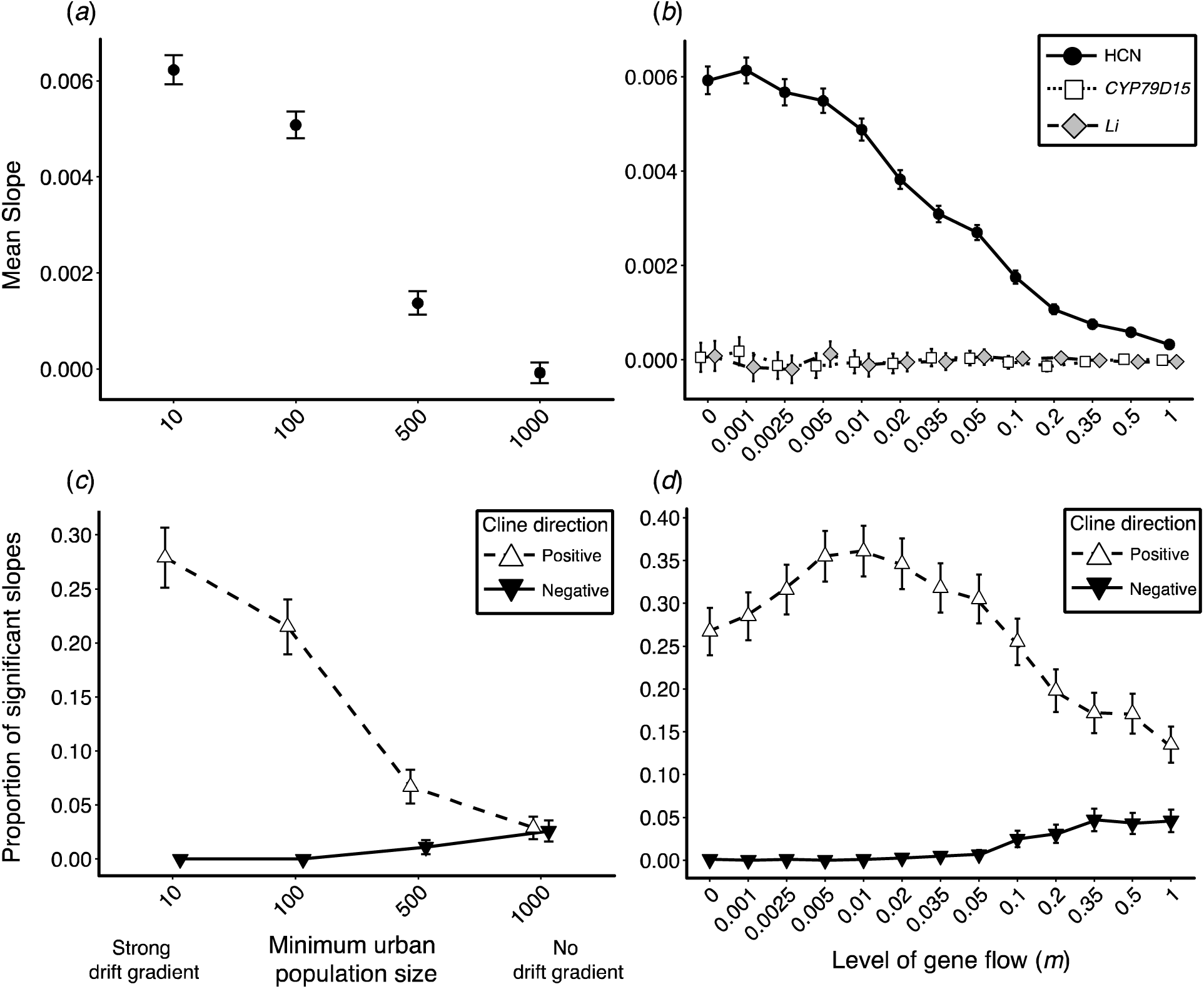
Spatial gradients in drift—controlled by varying the minimum urban population size across the landscape matrix (drift scenario 1)—influenced the formation of clines in HCN. (*a*) The mean strength of clines in HCN across 1000 simulations. (*b*) When there is a strong gradient in drift (minimum *urban N* = 10), gene flow influenced the mean strength of HCN (filled circles with solid line) clines, but not clines in *CYP79D15* (open squares with dotted line) or *Li* (grey diamond triangle with dashed line). (*c*) The proportion of significantly positive (open triangles with dashed line) and negative (black inverted triangles with solid line) clines under varying strengths of drift, simulated by varying the minimum urban population size. (*d*) When there is a strong gradient in drift (minimum *urban N* = 10), gene flow influences the proportion of significantly positive (open squares) and negative (filled diamonds) clines. All points represent mean or proportions ± 95% confidence intervals.

Gene flow reduced the mean slope of clines and the proportion of significantly positive clines. Under a strong spatial gradient in drift (urban *N* = 10), the strongest clines in the frequency of HCN (β ≈ 0.006) occurred with little to no gene flow, whereas increasing gene flow reduced the mean slope of clines to near zero (figure 2*b*). In contrast, the mean strength of clines at each of the two unlinked loci underlying HCN (i.e. *CYP79D15* and *Li*) was consistently zero for all levels of gene flow (figure 2b); at single loci (e.g. *CYP79D15* and *Li*) drift produces positive and negative clines in equal proportions, resulting in a mean slope of 0 when averaged across all clines (figure S12). Finally, the frequency of significantly positive clines peaked at ∼36 % when *m =* 0.01 and decreased to a minimum of ∼14% with increasing gene flow. Negative clines peaked at ∼4% at only the highest levels of gene flow (figure 2*d*).

### Question 2: How does selection affect the occurrence and strength of spatial clines in HCN?

In the absence of drift, selection influenced the formation of spatial clines in HCN. Independent of levels of gene flow, increasing the maximum strength of selection increased the mean strength of clines across 1000 simulations (figure 3*a*); selection increased the strength of clines from a slope of zero *s =* 0, to 0.037 when *s* = 0.2. Similarly, selection increased the frequency of significantly positive clines from 2.7% when *s* = 0 to 100% when *s* ≥ 0.025. Increasing gene flow reduced the effects of selection, leading to weaker clines for a given selection coefficient (figure 3*a*). Importantly, clines formed by selection (figure 3*a*) are consistently stronger than even the maximum strength of clines formed by drift, regardless of whether drift is manipulated by varying the maximum population size (figure 2*a*) or through serial founder effects (figure S3*a*).

**Figure 3:**
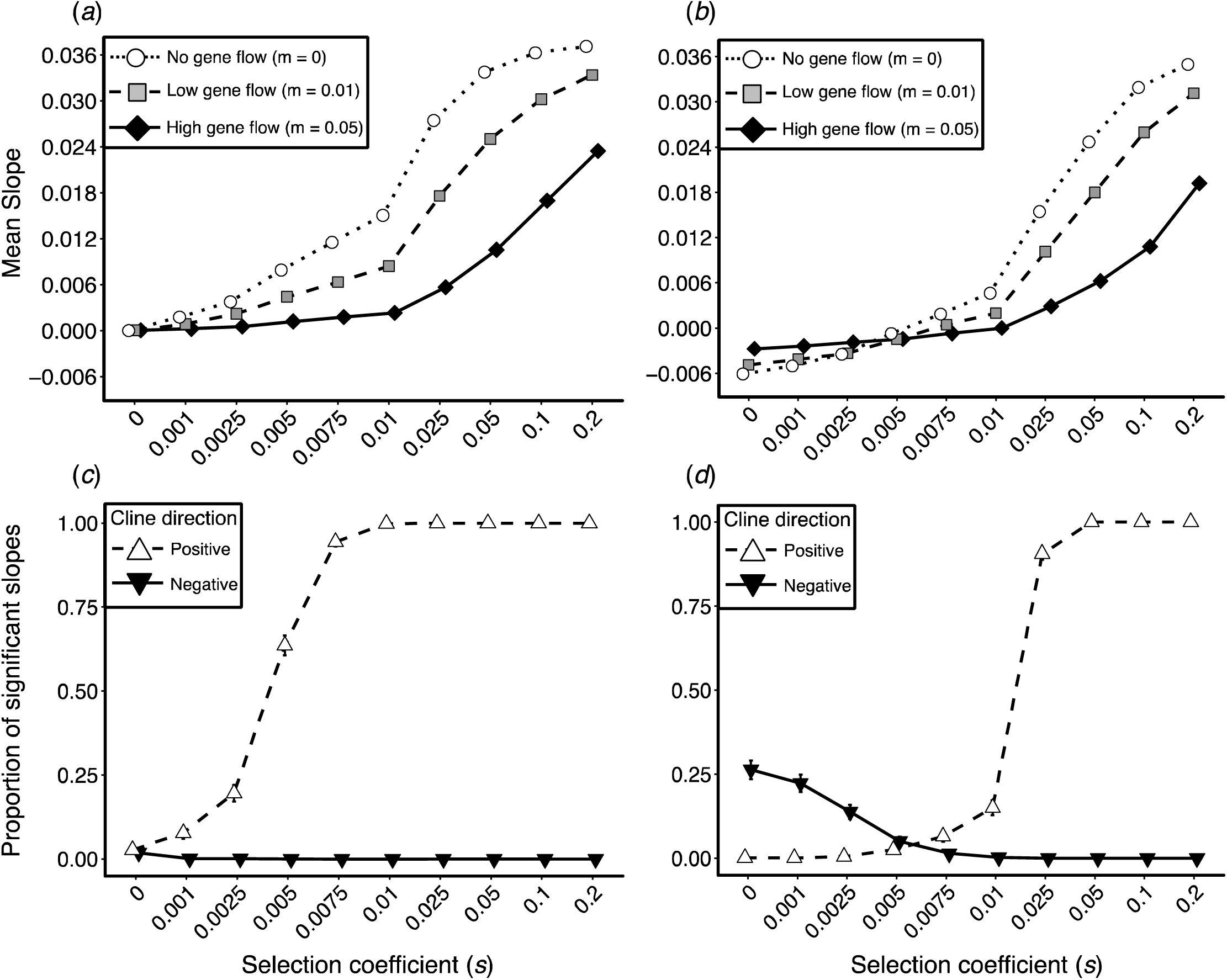
Selection influenced the formation of spatial clines in HCN in both the absence (*a* and *c*) and presence (*b* and *d*) of opposing gradients in drift. Selection favours HCN+ genotypes in rural populations and HCN– genotypes in urban populations. In (*b*) and (*d*), we imposed a spatial gradient in carrying capacity (i.e. drift scenario 1) such that the minimum *rural* population size was 10. As such, the stochastic loss of dominant alleles in smaller rural populations is countered by their higher fitness. In both the absence (*a*) and presence (*b*) of an opposing drift gradient, selection influenced the mean strength of clines across 1000 simulations under no (*m =* 0, open circles with dotted line), low (*m =* 0.01, grey squares with dashed line) and high (*m =* 0.05, black diamonds with solid line) gene flow, although this effect was reduced in the presence of drift. Similarly, selection influenced the proportion of significantly positive (open triangles with dashed lines) and negative (black inverted triangles with solid line) clines in both the absence (*c*) and presence (*d*) of an opposing gradient in drift. All points represent mean or proportions ± 95% confidence intervals.

### Question 3: What are the interactive effects of genetic drift and selection on the formation of clines in HCN?

In the presence of an opposing gradient in drift, selection generated fewer and weaker clines for all but the strongest selection coefficients. In the absence of gene flow, the mean slope of clines changed from negative to positive as the strength of selection increased. Clines were only consistently positive when the selection coefficient (*s*) was greater than 0.005 (figure 3*b* and 3*d*). By contrast, negative clines were more common when selection was less than 0.005, consistent with gradients in drift preferentially generating clines in HCN (see Question 1 above and figure 2*b* and 2*d*). Finally, drift increased the strength of selection necessary to produce 100% positive clines by 5-fold. That is, in the presence of strong drift (urban *N* = 10), 100% of clines were positive when *s =* 0.05 (figure 3*d*) whereas in the absence of drift, 100% of clines were positive when *s =* 0.01 (figure 3*c*).

## Discussion

Our simulations sought to understand the relative contributions of genetic drift, gene flow and selection for the evolution of parallel clines in hydrogen cyanide (HCN) along urbanization gradients. Several key results are most important in answering our research questions. We found that drift overwhelmingly led to the formation of positive clines in HCN, and stronger gradients in drift generated stronger clines, but these effects decreased with increasing gene flow (Question 1). Clines formed by drift alone were substantially weaker than those generated by selection, suggesting an upper limit to the strength of phenotypic clines in non-additive traits due to drift (Question 2). Finally, when selection operated counter to the prevailing gradient in drift, stronger selection was required to generate clines as strong as those observed in the absence of drift (Question 3). We begin by discussing the relevance of our results for the evolution of clines in cyanogenesis for white clover along urbanization gradients and other environmental gradients. We then suggest that urban environments—in combination with knowledge of the genetic architecture of focal traits—can provide the replication necessary for understanding the contributions of drift, gene flow and selection to the evolution of clines in non-additive traits. We end by more broadly discussing the role of drift and other evolutionary mechanisms in the formation of genetic and phenotypic clines.

### Evolution of cyanogenesis clines in white clover

Understanding geographical variation in allele and phenotype frequencies often provides insight into the evolutionary mechanisms structuring patterns of genetic variation in natural populations. In white clover, pioneering work by Hunor Daday identified broad-scale latitudinal clines in the frequency of HCN across multiple continents [30, 31], and altitudinal clines across the central European Alps [32]. Subsequent work has shown that clines in the frequency of HCN are common, with higher frequencies of HCN occurring in warmer and drier habitats [33–37]. These patterns are thought to reflect the benefits of producing HCN in warmer environments where herbivore damage is greater [36, 38], the benefits of producing cyanogenic glycosides in regions subject to moderate drought stress [36, 37], and the cost of producing HCN in frost-prone habitats [36, 39]. Consistent with this latter view, urban-rural clines in HCN appear to be driven by lower urban winter ground temperatures acting as a selective agent against cyanogenic plants, resulting in lower HCN frequencies in urban populations [23]. Nonetheless, investigations into the mechanisms structuring spatial variation in HCN frequencies have focused largely on selection (but see [26, 35]), despite the genetic architecture of HCN making populations particularly susceptible to loss of HCN via drift.

The presence of repeated correlations between environmental variables and phenotype frequencies is often considered strong evidence for the role of natural selection in generating adaptation. However, clines may also form via neutral processes [7, 8] and this may be exacerbated in traits with an epistatic genetic architecture. For example, our simulations showed that gradients in the strength of drift from rural (weak drift) to urban (strong drift) populations preferentially led to urban populations having lower HCN frequencies, consistent in direction with phenotypic clines in HCN observed across cities [23, 26]. Thus, when a trait’s genetic architecture predisposes populations to evolve in a particular direction under the effects of drift alone, drift should be rejected as a null hypothesis prior to invoking selection’s role in generating clines. Observing more clines in natural populations than would be expected under drift alone is one way to address this problem. For example, null distributions for the proportion of clines expected under drift can be generated through simulations and compared to the actual proportion observed in nature. Ideally, simulations would be parameterized with empirical estimates of gene flow and effective population sizes to provide system-specific estimates of expected proportions. Unfortunately, we lack these data for white clover populations making it difficult to compare the clines observed across cities to the proportions estimated in these simulations.

While drift preferentially generated positive clines in HCN, the mean strength of clines formed by drift was substantially lower than those involving selection. A possible explanation for this is that although strong drift in urban populations drives HCN frequencies downward, the weak drift in rural populations and the absence of selection results in rural HCN frequencies fluctuating around their initial frequency. By contrast, simulations involving selection favoured HCN+ genotypes in rural populations and HCN– genotypes in urban populations, resulting in a steeper gradient in phenotype frequencies between urban and rural populations and thus stronger clines. To understand whether drift is likely to generate clines in HCN as strong as those observed across cities, we compared the magnitude of standardized slopes from our simulations (text S7, figure S13) to those observed across cities for which there are data [23, 26]. Even under an intermediate gradient in drift (urban *N* = 100), high gene flow (*m* = 0.05) and no selection there was substantial overlap in the distribution of standardized slopes from observed and simulated data (–0.30 < β_simulated_ < 0.27, figure S13*d*; –0.08 < β_observed_ < 0.30, figure S13*e*), suggesting drift can generate clines as strong as those observed across cities. However, recent empirical work found no decrease in neutral genetic diversity in urban populations [26] and identified clines in the dominant alleles at individual loci underlying cyanogenesis [23], suggesting that genetic drift may not be an important mechanism structuring urban-rural phenotypic clines in HCN [26]. Similarly, the absence of population structure and isolation by distance at neutral microsatellite markers in North American clover populations suggests that latitudinal clines in HCN are adaptive [35], at least in the invaded range. Thus, while there is overlap in the strength of observed clines and those simulated by drift, current evidence from neutral markers strongly suggests that many clines in cyanogenesis are adaptive.

### Urban environments as replicated systems to study evolution

Clines have traditionally been studied along latitudinal [31,40,41], altitudinal [32,34,42] and longitudinal [10, 37] transects, because such transects are often associated with gradients in putative selective agents (e.g. temperature, precipitation, biotic interaction strength). Importantly, geographical transects can also be associated with variation in demographic factors important for determining the extent to which drift will act in populations. This may be especially true during invasions or rapid range expansions when founder events result in smaller marginal populations that exchange fewer migrants, leading to reduced genetic diversity and increased differentiation relative to more central populations [24, 43]. Such spatial variation in demography may provide the foundation upon which parallel clines in non-additive traits can form. Studies of non-additive trait differentiation across replicate geographical gradients in selection and demography, coupled with knowledge of the focal trait’s genetic architecture, would provide a strong test of the relative contribution of drift, gene flow and selection in the formation of parallel clines. However, logistical challenges associated with replicating continental-scale latitudinal, altitudinal or longitudinal transects has precluded such replication in most systems.

Urban environments represent globally replicated disturbances to the landscape and are frequently associated with adaptive and non-adaptive evolutionary changes within populations [44]. The frequent observation of increased drift in urban populations, together with our results showing parallel changes in HCN frequencies under gradients of drift, suggests that urban populations may be prone to directional phenotypic change due solely to stochastic forces. The importance of drift in structuring urban phenotype frequencies will vary across cities and depend on the extent to which urban fragmentation has reduced the availability of suitable habitat for the focal species and whether sufficient connectivity exists to enable gene flow between populations. For example, urban construction and fragmentation is predicted to reduce local populations sizes (e.g. [45]) but corridors that facilitate gene flow (e.g. [46]) may limit drift’s ability to generate genetic and non-additive trait differentiation between populations.

The natural history and biology of focal taxa is also likely to affect their susceptibility to stochastic changes in allele and phenotype frequencies. Owing to their tolerance of disturbed habitats, non-native species often increase in frequency in urban environments at the expense of native species, which are more susceptible to environmental changes associated with the development of cities [47]. Native species (i.e. those that pre-date urbanization) may therefore be more vulnerable to stochastic changes in demography with increasing urbanization leading to increased genetic drift in urban populations of native, but not exotic, species. In support of this view, red-backed salamanders [22], white-footed mice [21] and fire salamanders [45]—three species native to the regions in which they were studied—all show reductions in effective population size or reduced neutral genetic diversity in urban populations. By contrast, white clover, originally native to Eurasia, shows inconsistent effects of urbanization on neutral genetic diversity across 8 cities in Ontario, Canada [26], suggesting no increased effects of drift in urban populations. In this very limited comparison, the data suggest that urban populations of native species appear more susceptible to the effects of genetic drift than non-native species. Predictions on the effects of urbanization on the strength of genetic drift should therefore be based at least in part on the natural history of species being studied.

### Genetic drift and the formation of clines

The ability for gradients in the strength of genetic drift to generate clines in phenotypic traits depends largely on the genetic architecture of the trait being examined. For example, drift operating alone should generate clines in additive quantitative traits or at individual loci in proportion to the initial allele frequencies. In such cases, there will typically be an equal proportion of positive and negative clines with no consistent change in allele or phenotype frequency when averaged across all simulated clines [8]. Our results support this prediction: in the absence of gene flow and selection, clines at individual loci underlying HCN (i.e. *CYP79D15* and *Li*) occurred with equal frequency resulting in a mean slope of zero across 1000 simulations (figure 2*b*, figure S12). By contrast, clines in HCN were overwhelmingly positive because the epistatic interaction among underlying loci makes natural populations particularly susceptible to the formation of clines via drift. This prediction can provide a way of assessing whether observed clines are due to drift or other evolutionary mechanisms; repeated clines in the same direction at individual loci underlying non-additive traits excludes drift as a possible mechanism producing clines.

While genetic drift is sufficient to generate clines in non-additive traits, it is insufficient on its own to maintain them. At equilibrium, genetic drift is expected to fix different alleles in different populations [48], leading to the eventual elimination of clines. Our results support this as in the absence of migration, clines formed by drift became gradually weaker over time (text S6, figure S11). However, small amounts of gene flow, which averages allele frequencies among neighboring populations and slows the loss or fixation of alleles [48] can allow for more long-term maintenance of clines. Selection, on the other hand, can maintain clines indefinitely when the alternate phenotypes are each favoured on either end of the cline, although clines will become increasingly non-linear as time proceeds and populations become fixed for alleles on opposing sides of the population where phenotypes have equal fitness. Again, gene flow may slow the fixation of alleles resulting in more long-term maintenance of smooth clines in phenotype frequencies [5].

The presence of gradients in the strength of drift has important consequences for the ability of selection to generate clines in non-additive traits. We found that gradients in drift that oppose selection can constrain the formation of parallel clines due to selection. The presence and strength of clines in such cases thus reflects a balance between drift and selection: drift may lead to directional changes in the frequency of non-additive traits, but this may be countered by selection if the fitness differential between genotypes is sufficiently strong. An interesting scenario not modeled here concerns the ability of selection to maintain clines formed via drift. Using HCN as an example, if selection is constant and favours HCN+ genotypes across the entire transect, the increased fitness of HCN+ genotypes may counter their stochastic loss in urban populations via drift. Equilibrium could be reached when the rate at which the frequency of a non-additive trait decreases via drift equals the rate at which its frequency increases due to selection. The maintenance of clines in non-additive traits under drift-selection equilibrium represents an interesting avenue for future research.

The formation of phenotypic clines in non-additive traits via neutral processes is likely a common phenomenon. We have shown that gradients in the strength of drift lead to deterministic phenotypic clines in HCN. This, together with predictable changes in other epistatically determined phenotypes due to stochastic processes (e.g. *Eichhorinia paniculata*, [12]), suggests that non-additive traits are especially susceptible to deterministic changes in frequency via stochastic forces. When a phenotype is the end-product of a multigene pathway, any loss of function mutation to a protein in the pathway will knock-out the phenotype. Because many loss-of-function mutations can be complemented in diploids, the genetic architecture of HCN (i.e. duplicate recessive epistasis) is likely common. Further, the more genes in the pathways, the higher the probability of generating the knock-out phenotype by drift. Additional theoretical and empirical work exploring changes in the frequency of non-additive phenotypes—whether due to epistasis or dominance at individual loci—is required to generate predictions and assess the generality of this phenomenon in natural populations.

## Conclusion

We have shown that gradients in the strength of drift can lead to repeated spatial clines in the frequency of a non-additive trait, despite equal frequencies of positive and negative clines at underlying loci. Drift should thus be considered a null model to be rejected prior to invoking the selection in the formation of clines, especially when phenotypes result from interactions among multiple genes or metabolic pathways. Rejecting drift as a mechanism producing clines in non-additive traits can come in a number of ways. First, showing the absence of clines at neutral loci across the genome despite the presence of clines in the focal trait is strong evidence that adaptive processes are at work as drift is expected to affect all loci. Second, showing that more clines or stronger clines are observed in nature than would be expected under drift suggests that non-neutral mechanisms are responsible for generating clines. A disadvantage of this approach is that it does not inform the mechanism structuring any one cline but rather rejects drift as a mechanism producing all clines. Third, showing that individual loci underlying focal traits consistently form clines in the same direction strongly suggests that other mechanisms are generating clines since drift should not display directionality at individual loci. The second and third points above require large-scale replication and we suggest that urban environments can provide the replication necessary to understand the relative contributions of adaptive and non-adaptive processes in the formation of parallel clines. Finally, we caution that observations of parallel clines may represent modern “spandrels” [48]; studies presuming that repeated phenotypic clines are evidence of adaptation must take explicit consideration of the genetic architecture underlying the focal trait to model the expected change in phenotypic frequency from non-adaptive processes.

## Data and code accessibility

Data available from the Dryad Digital Repository: https://doi.org/10.5061/dryad.6nv2t4p. All Python and R code used in generating and analyzing data can be found on the GitHub page for J.S.S. (https://github.com/James-S-Santangelo/Simulating-Evolutionary-Clines-SEC-).

## Author contributions

R.W.N., M.T.J.J. and J.S.S. conceived of the study. J.S.S. designed the simulations with input from R.W.N. and M.T.J.J. J.S.S. analyzed the data and wrote the first draft of the manuscript with all authors contributing to subsequent drafts.

## Acknowledgements

We thank B. Cohan, A. Dao, C. Fitzpatrick, M. Hetherington-Rauth, S. Innes, V. Nhan, R. Rivkin, A. Schneider and two anonymous reviewers for providing helpful comments that improved the manuscript. HPCNODE1 and Brian Novogradac provided the computational resources necessary to run the simulations.

