## Supplementary Materials for "Modern spandrels: The roles of genetic drift, gene flow and natural selection in the formation of parallel clines"

**Supplementary figures**

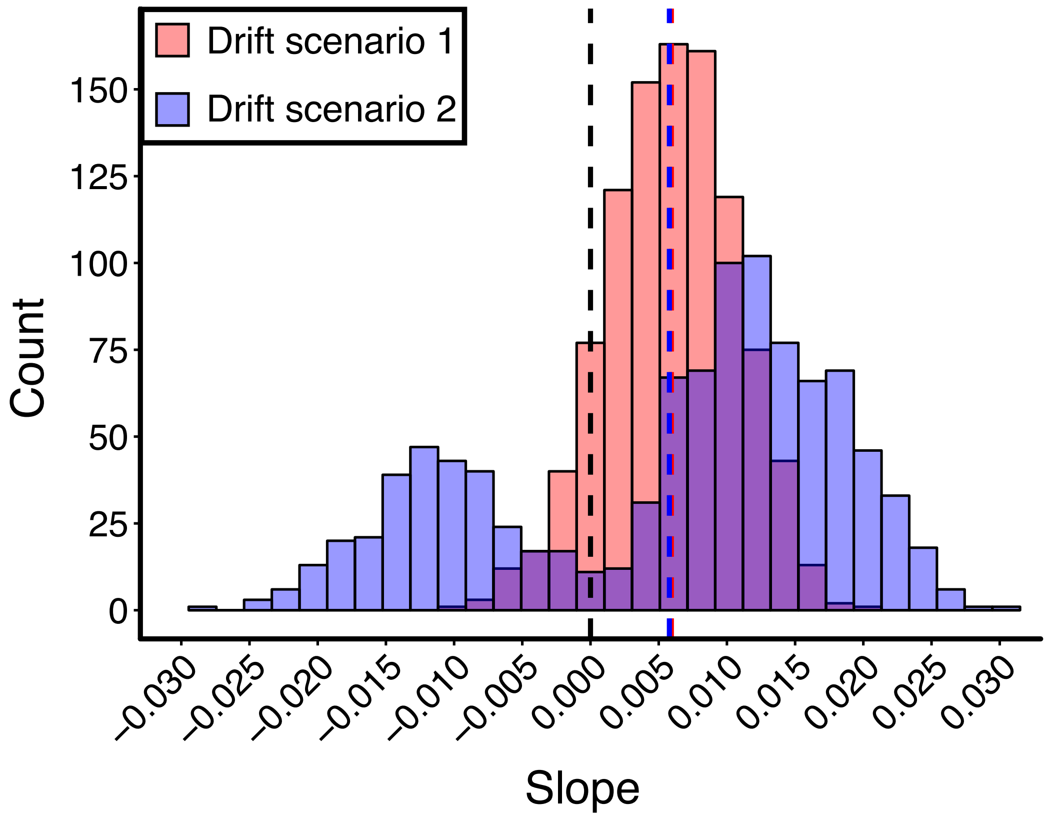


**Figure S1:** Distribution of slopes from 1000 simulations for drift scenario 1 (i.e. gradient in carrying capacity, red) and drift scenario 2 (i.e. serial founder events, blue). Slopes from both scenarios are plotted based on the parameter combinations that produced the strongest clines: slopes from drift scenario 1 are from a strong gradient in drift (i.e. rural $N = 1000$, urban $N = 10$) while those from drift scenario 2 are from simulations involving intermediate founding proportions (proportion = 0.2). Slopes from both scenarios were simulated in the absence of gene flow (i.e. $m = 0$). Black dashed line represents a slope of zero. While drift scenario 2 generated more negative clines and clines of more extreme magnitude, the distributions are largely overlapping and there is little difference in mean slope across all 1000 simulations for both scenarios (red and blue dashed lines).Thus, independent of drift scenario, our key result remains the same: gradients in the strength of drift lead to parallel clines in the frequency of HCN.

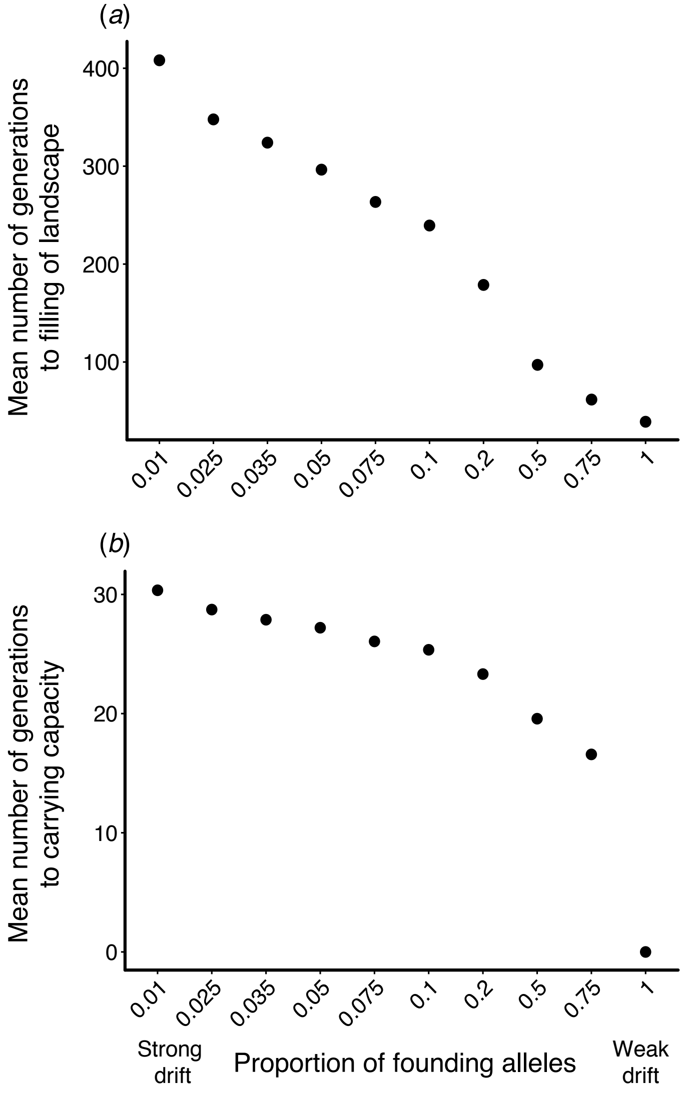


**Figure S2:** Serial founder events (i.e. drift scenario 2) influenced (*a*) the number of generations require for the linear landscape to fill with populations and (*b*) the number of generations required for the urban-most population to reach carrying capacity ($K = 1000$) once it has been founded. Points in (*a*) and (*b*) represent the mean number of generations across 1000 simulations. 95% confidence intervals are plotted but are hidden by points due to minimal variation in the number of generations between simulations.

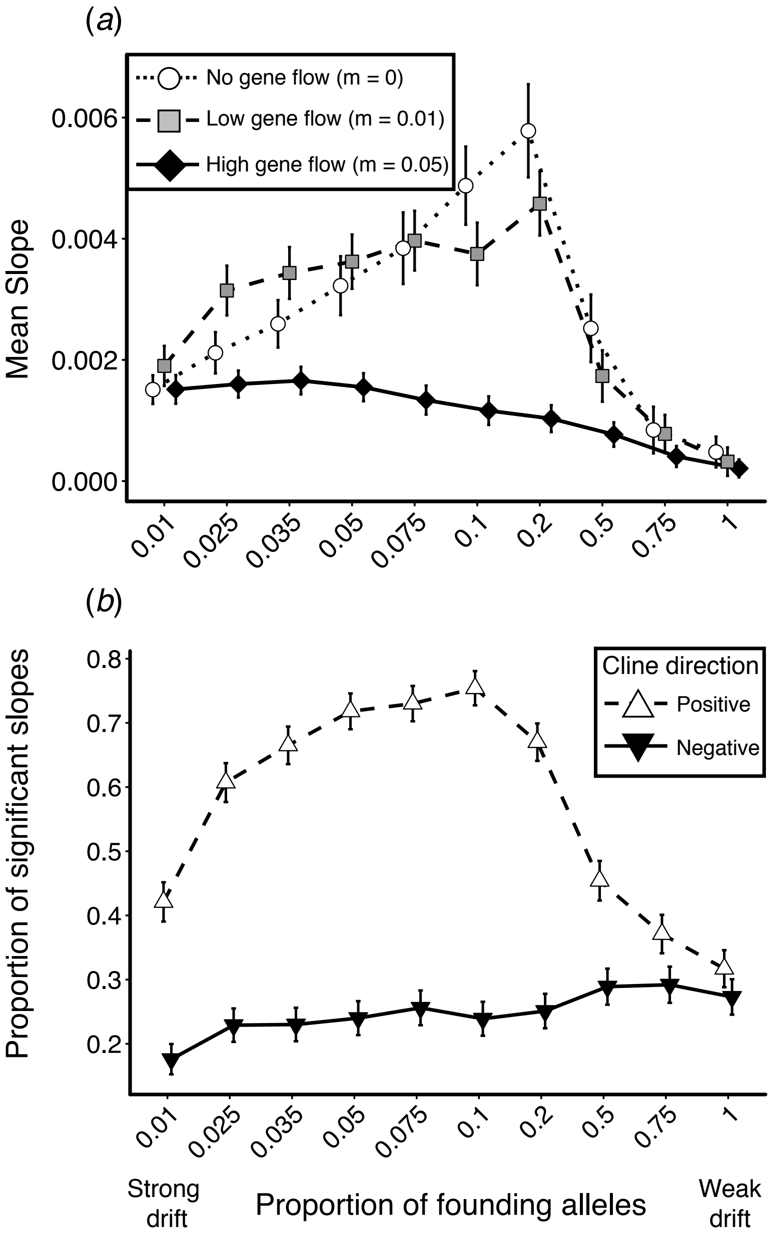


**Figure S3:** Serial founder events (i.e. drift scenario 2) influenced the mean strength of clines and the proportion of significatly positive and negative clines. The strength of founder events is represented as the proportion of alleles sampled to form the newly colonized population, which is equivalent to sampling a finite number of individuals. (*a*) Shown are the effects of serial founder effects on the mean strength of clines across 1000 simulations under three gene flow rates: no gene flow ($m$ = 0, open circles with dotted line), low gene flow ($m$ = 0.01, grey squares with dashed line) and high gene flow ($m$ = 0.05, black diamonds with solid line). (*b*) Serial founder events influence both the proportion of significantly positive (open triangles with dashed line) and negative (black inverted triangles with solid line) clines. All points represent mean or proportions ± 95% confidence intervals.

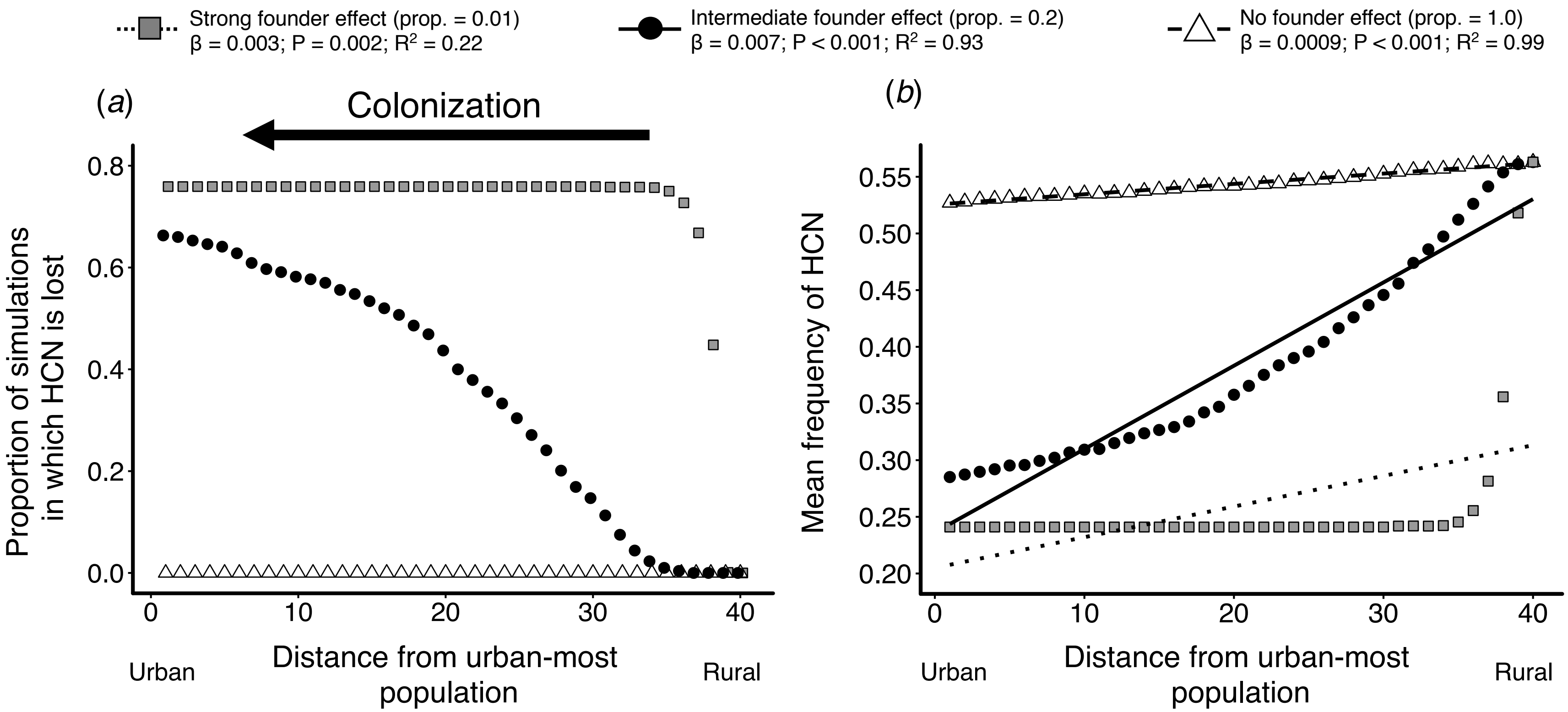


**Figure S4:** The strength of founder events (i.e. drift scenario 2) influenced the pace at which HCN was lost from populations during colonization and ultimately the strength of phenotypic clines in HCN. (*a*) Proportion of 1000 simulations where HCN is lost (i.e. frequency = 0) for each population in the landscape (i.e.1 to 40) under strong founder effects (proportion of founding alleles = 0.01, grey squares), intermediate founder effects (proportion = 0.2, black circles) and no founder effects (proportion = 1.0, open triangles). (*b*) Linear regressions of mean within-population HCN frequency across 1000 simulations against a population's position in the landscape matrix for strong founder effects (grey squares with dotted regression line), no founder effects (open triangle with dashed regression line), and intermediate founder effects (black circle with solid regression line). β coefficients, *P*-values and $R^2$ in legend refer to regressions run in (*b*).

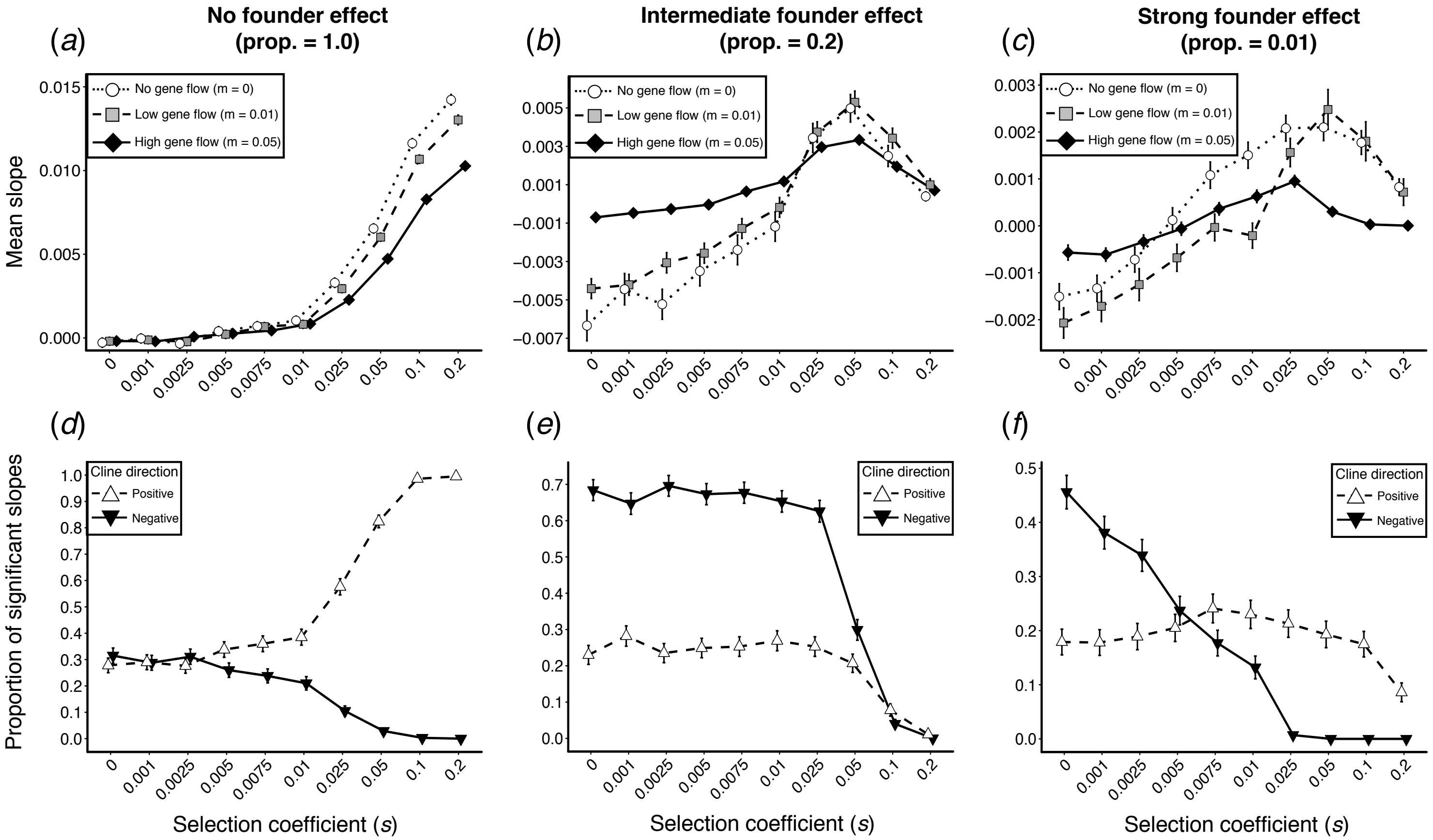


**Figure S5:** Serial founder events (i.e. drift scenario 2) and selection interacted in the formation of spatial clines in HCN. Populations colonized from the urban-most population to the rural-most population. Selection favoured HCN+ genotypes in rural populations and HCN– genotypes in urban populations. (*a* – *c*) The mean slope of clines in HCN across 1000 simulations under no gene flow (open circles with dotted line), low gene flow (grey square with dashed line), or high gene flow (black diamonds with solid line). (*d* – f) The proportion of significantly positive (open triangles with dashed line) and negative (black inverted triangle with solid line) clines across 1000 simulations. Founder events represent the proportion of alleles sampled in founding the new population and are either absent (proportion = 1.0, *a* and *d*), of intermediate strength (proportion = 0.2, *b* and *e*) or strong (proportion = 0.01, *c* and *f*). All points represent means or proportion ± 95% confidence intervals.

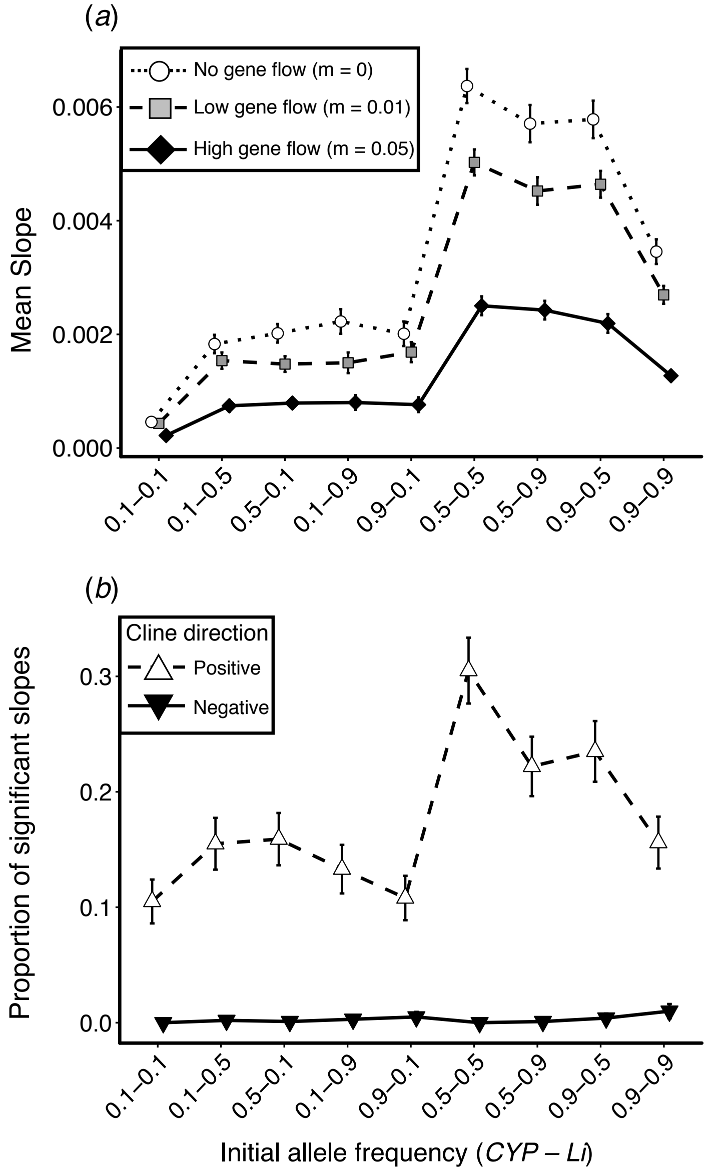


**Figure S6:** Effects of initial frequency of both dominant alleles (*CYP79D15* and *Li*) on (*a*) the mean strength of clines across 1000 simulations and (*b*) the proportion of significantly positive (open triangles with dashed line) and negative (black inverted triangles with solid line) clines. Simulations were run under a strong gradient in drift (i.e. drift scenario 1), manipulated by imposing a gradient in the maximum size of populations: rural populations were large ($N$ = 1000) while urban population were small ($N$ = 10). In (*a*) we examined the mean slope of clines under no (open circles with dotted line), low (grey square, with dashed line), and high (black diamonds with solid line) gene flow. In (*b*) positive clines reflect significantly ($P < 0.05$) less HCN in urban populations relative to rural populations while negative clines reflect the opposite. All points represent means or proportions ± 95% confidence intervals.

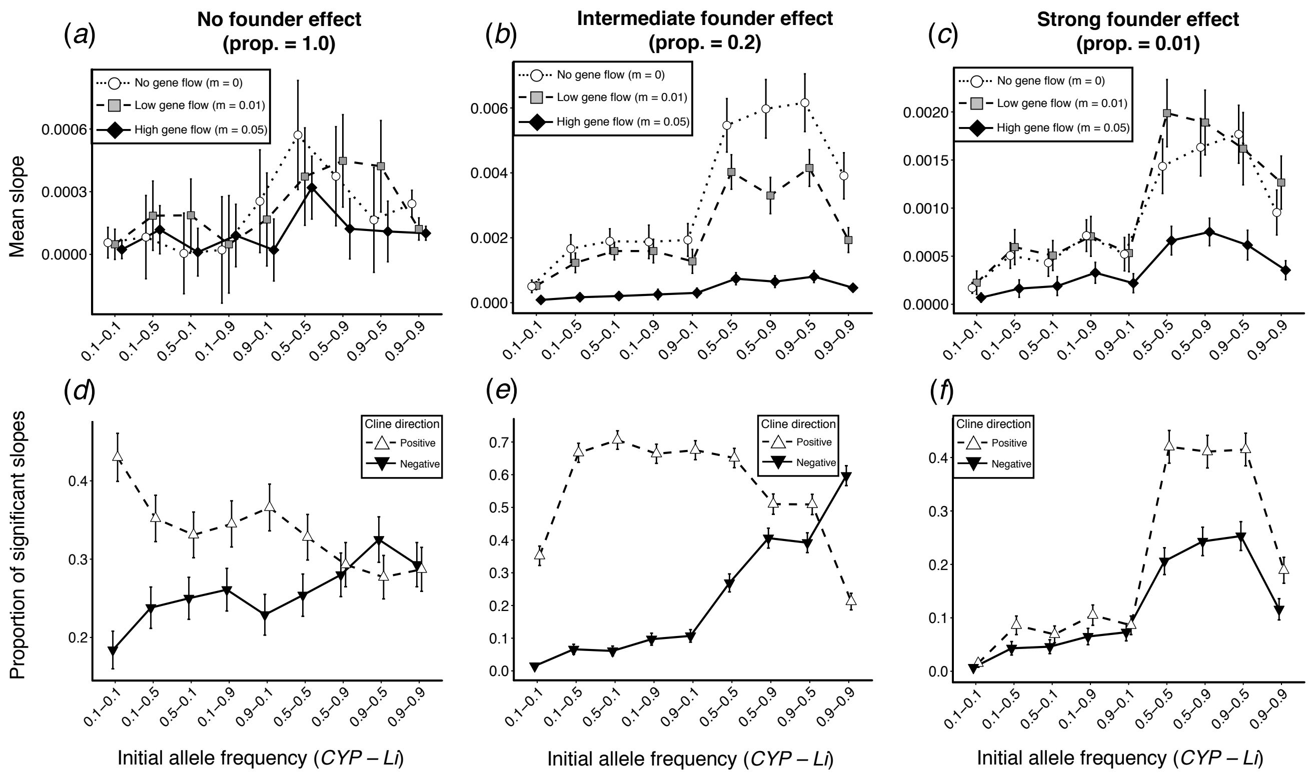


**Figure S7:** Serial founder events (i.e. drift scenario 2) and the initial frequency of the dominant alleles interacted in the formation of spatial clines in HCN. Populations colonized from the urban-most population to the rural-most population. (*a* – *c*) The mean slope of clines in HCN across 1000 simulations under no gene flow (open circles with dotted line), low gene flow (grey square with dashed line), or high gene flow (black diamonds with solid line). (*d* – f) The proportion of significantly positive (open triangles with dashed line) and negative (black inverted triangle with solid line) clines across 1000 simulations. Founder events represent the proportion of alleles sampled in founding the new population and are either absent (proportion = 1.0, *a* and *d*), of intermediate strength (proportion = 0.2, *b* and *e*) or strong (proportion = 0.01, *c* and *f*). All points represent means or proportion ± 95% confidence intervals.

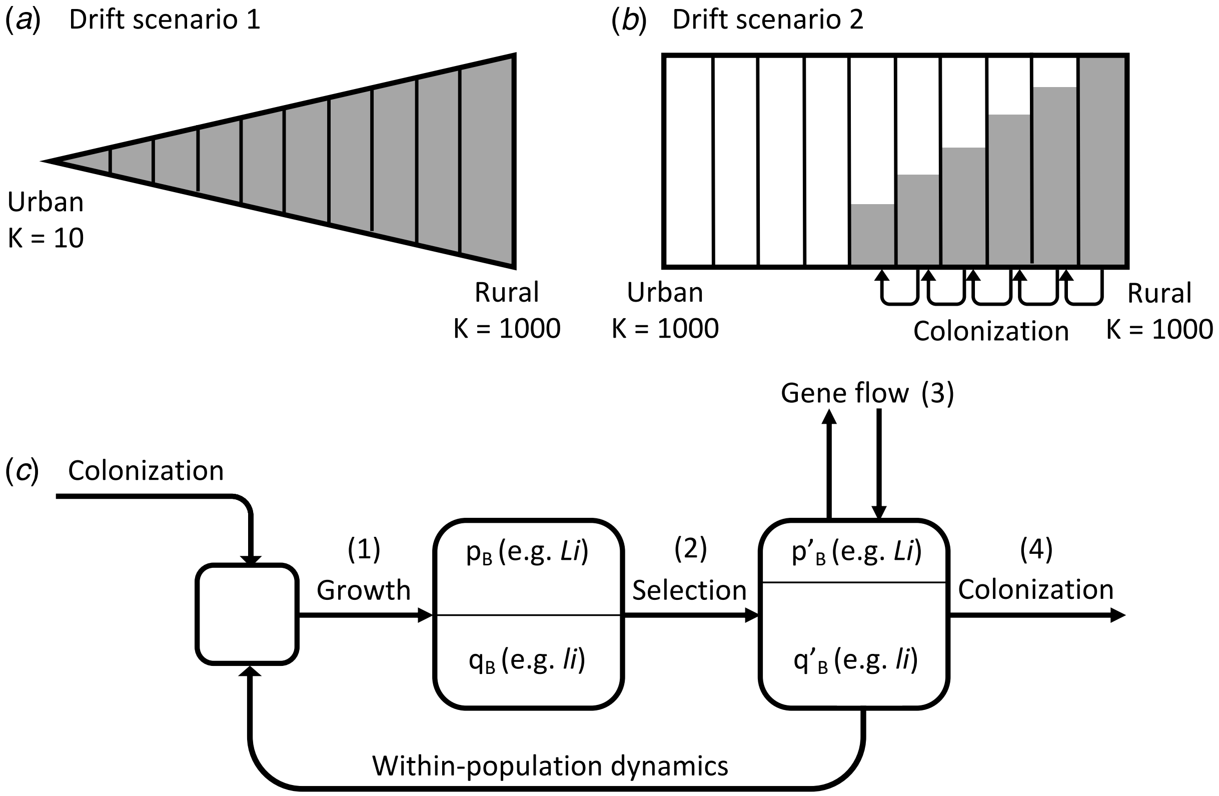


**Figure S8:** Diagrammatic representation of simulations examining the effects of genetic drift, gene flow and selection on spatial clines in HCN. We manipulated the effects of drift in two ways: (*a*) In drift scenario 1 a spatial gradient in carrying capacity (*K*) was created across the linear matrix, which placed an upper limit on the size (*N*) of each population. For most simulations (see Table S2), population size was greatest in the rural-most population ($N = 1000$) and declined linearly to the urban-most population ($N = 10$). In this scenario, all patches (separated by solid vertical lines) started with populations at their carrying capacity in generation one (represented by grey filling of patches). (*b*) In drift scenario 2, the strength of drift was manipulated through serial founder events during the colonization of the urban environment, beginning with a single rural population at carrying capacity. Populations could only colonize adjacent patches and the proportion of founding alleles was varied to control the strength of drift (i.e. lower proportion = stronger drift). (*c*) Schematic of the order of events during simulations of drift scenario 2 (i.e. *b*, numbers represent order of events). Boxes represent a single population as it proceeds through the simulations. Upon colonization, populations first grow according to a logistic growth model (growth rate [r] = 1.5). Populations are then subject to selection, followed by gene flow and finally colonization of adjacent patches. Each generation, we tracked the frequency of dominant alleles at both loci underlying HCN production (i.e. *CYP79D15* and *Li*) and the frequency of HCN within each population in the matrix.

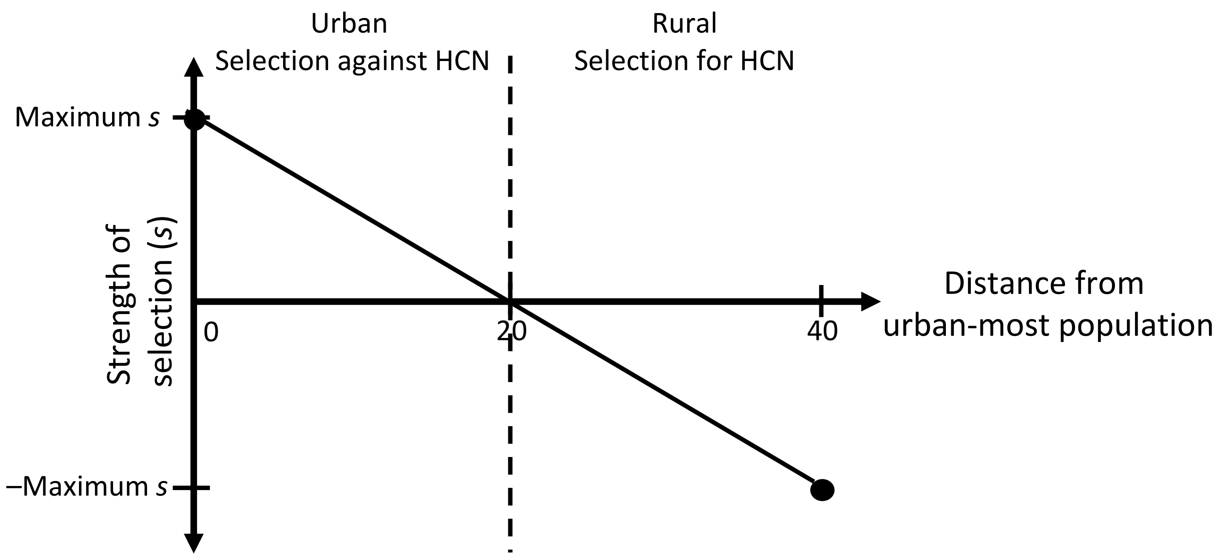


**Figure S9:** The strength of selecting favoring cyanogenic (i.e. HCN+) or acyanogenic (i.e. HCN–) genotypes depended on the population's position on the landscape. We first defined a maximum selection coefficient (-*s* to *s*), which favoured HCN– genotypes in urban populations and HCN+ genotypes in rural populations. The selection coefficient varied linearly across the matrix such that HCN+ and HCN– genotypes had equal fitness in the central population of the landscape (i.e. population 20).

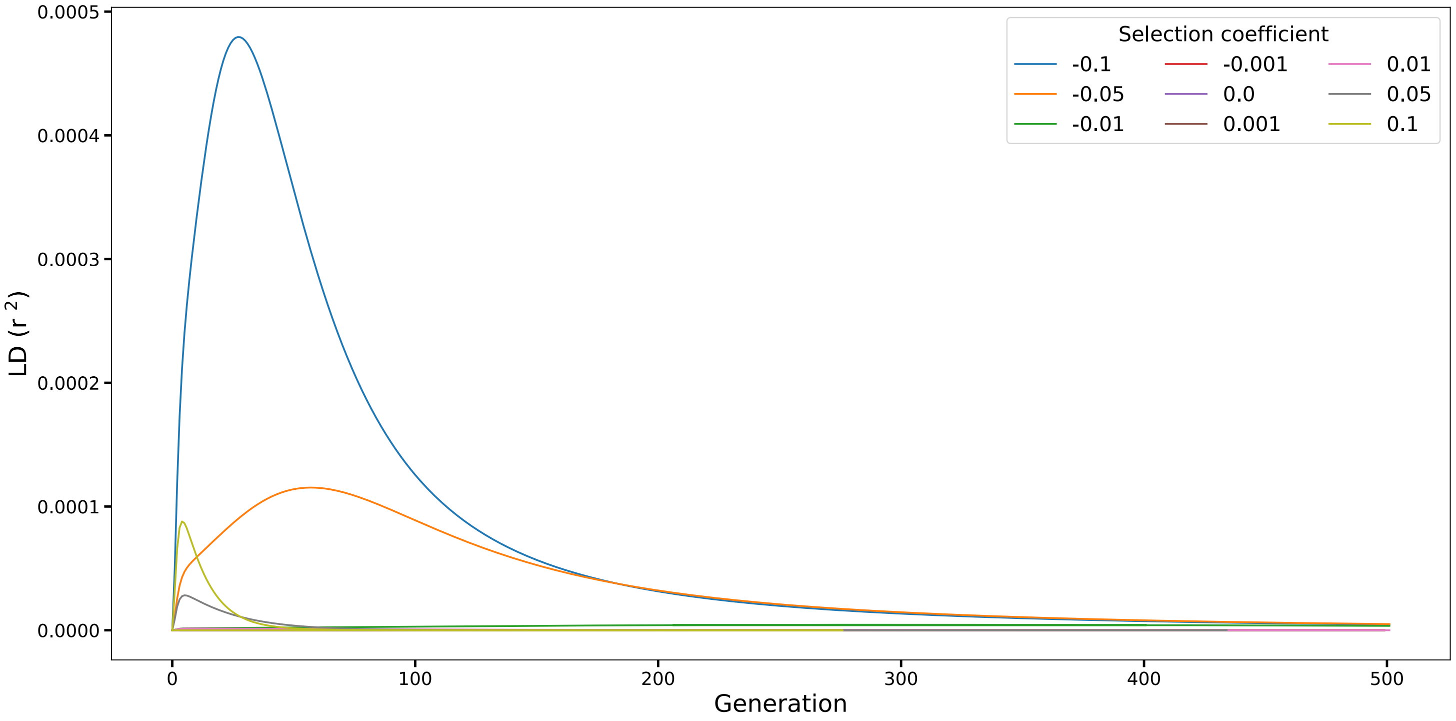


**Figure S10:** The build-up of linkage disequilibrium (LD) between *CYP79D15* and *Li* due to selection acting on cyanogenic white clover genotypes. Negative selection coefficients represent selection acting against cyanogenic clover genotypes while positive coefficients represent selection favouring cyanogenesis. Selection causes minimal build-up of LD between *CYP79D15* and *Li*, which decays rapidly over 500 generations.

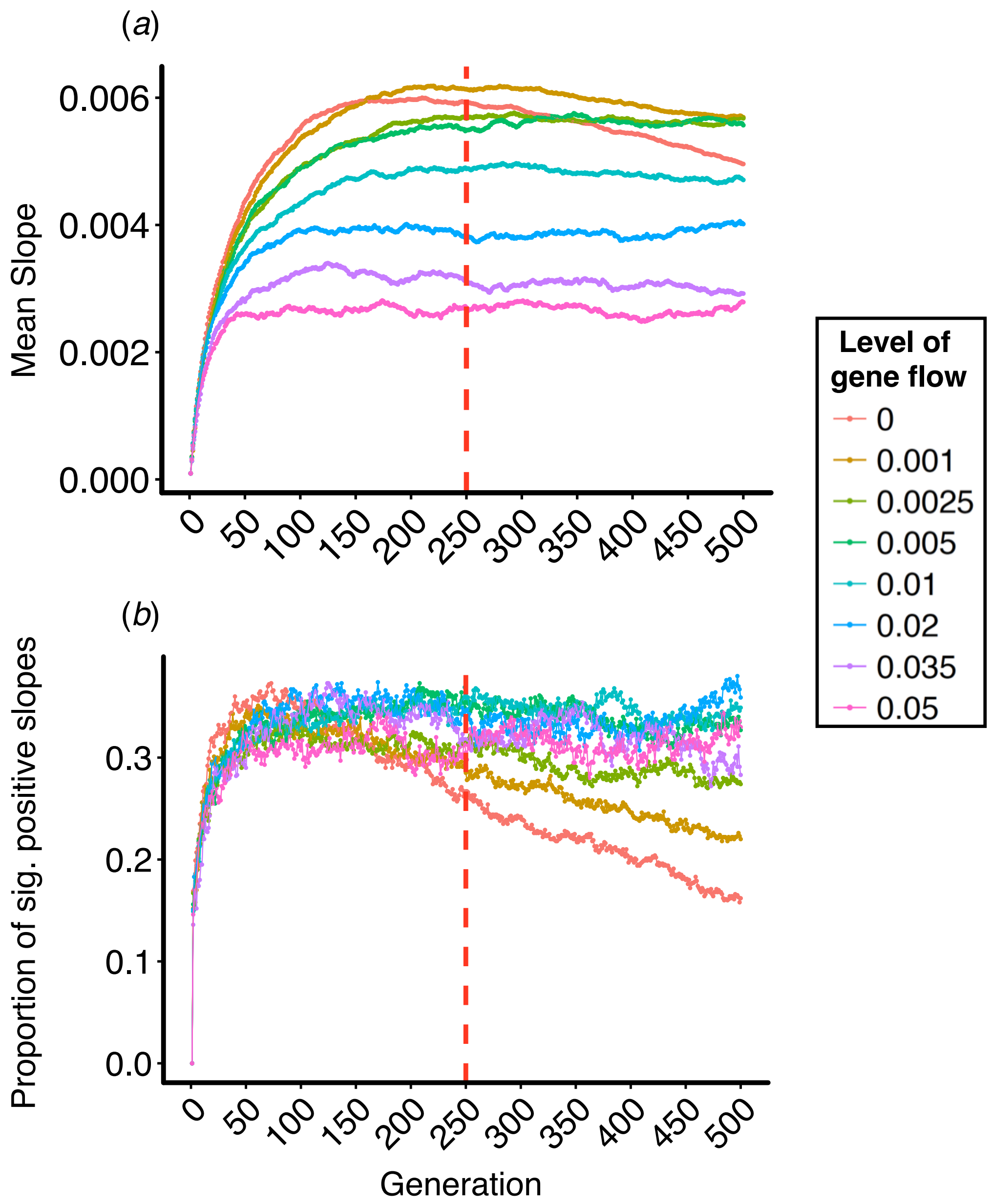


**Figure S11:** Differences in (*a*) the mean strength of clines and (*b*) the proportion of significantly positive clines remain qualitatively similar regardless of which generation is chosen for analysis. (*a*) The mean strength of clines across 1000 simulations every generation from 1 to 500 under varying levels of gene flow. (*b*) The proportion of significantly positive clines across 1000 simulations every generation from 1 to 500 under varying levels of gene flow. Only levels of gene flow from 0 to 0.05 are shown to increase visibility of lines. Red bar at generation 250 represents the generation used in analyses and the results are shown in main text (figures 2*b* and 2*d*).

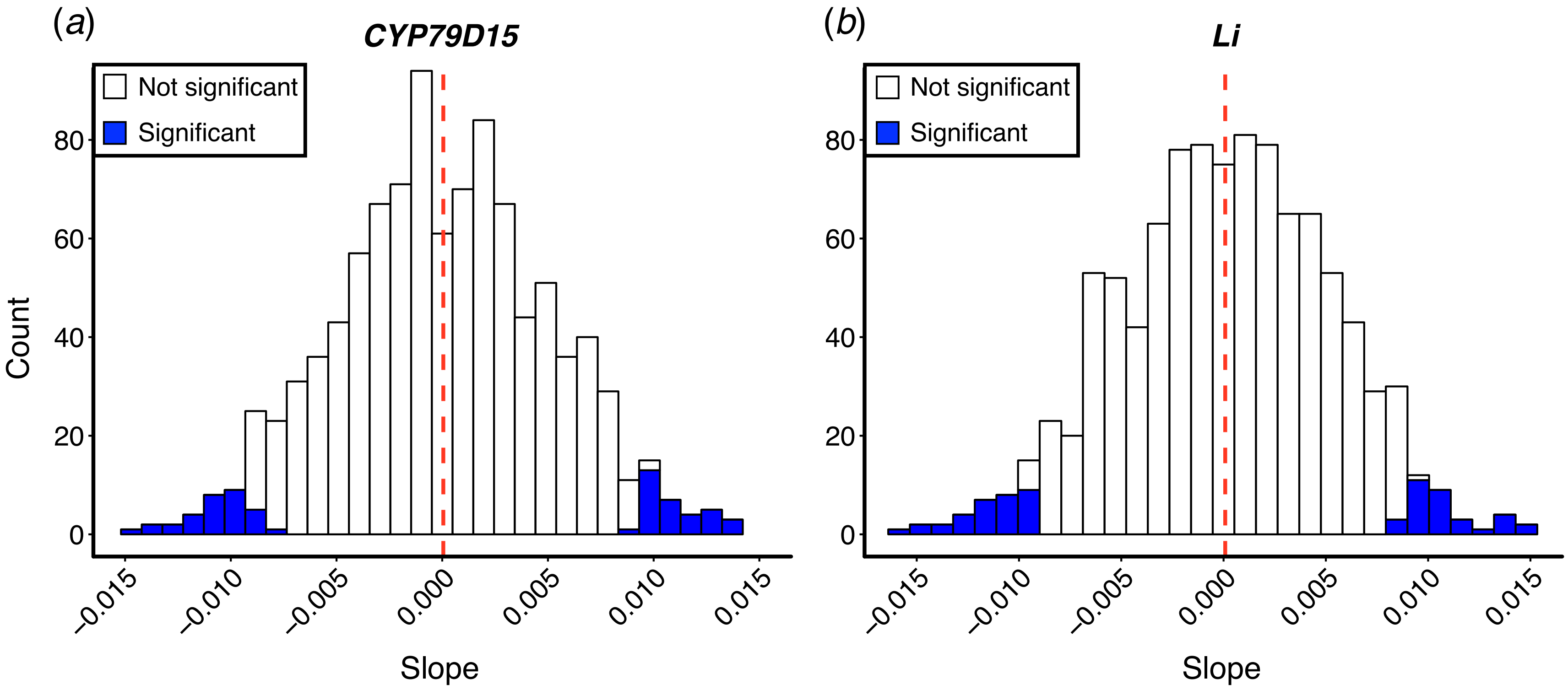


**Figure S12:** Distribution of slopes for the (*a*) *CYP79D15* locus and (*b*) *Li* locus underlying the production of HCN. Blue bars represent clines that are significant at $P < 0.05$. Red dashed bar represents the mean slope. We show only the distribution of slopes in the absence of gene flow, but results are qualitatively similar for all levels of gene flow.

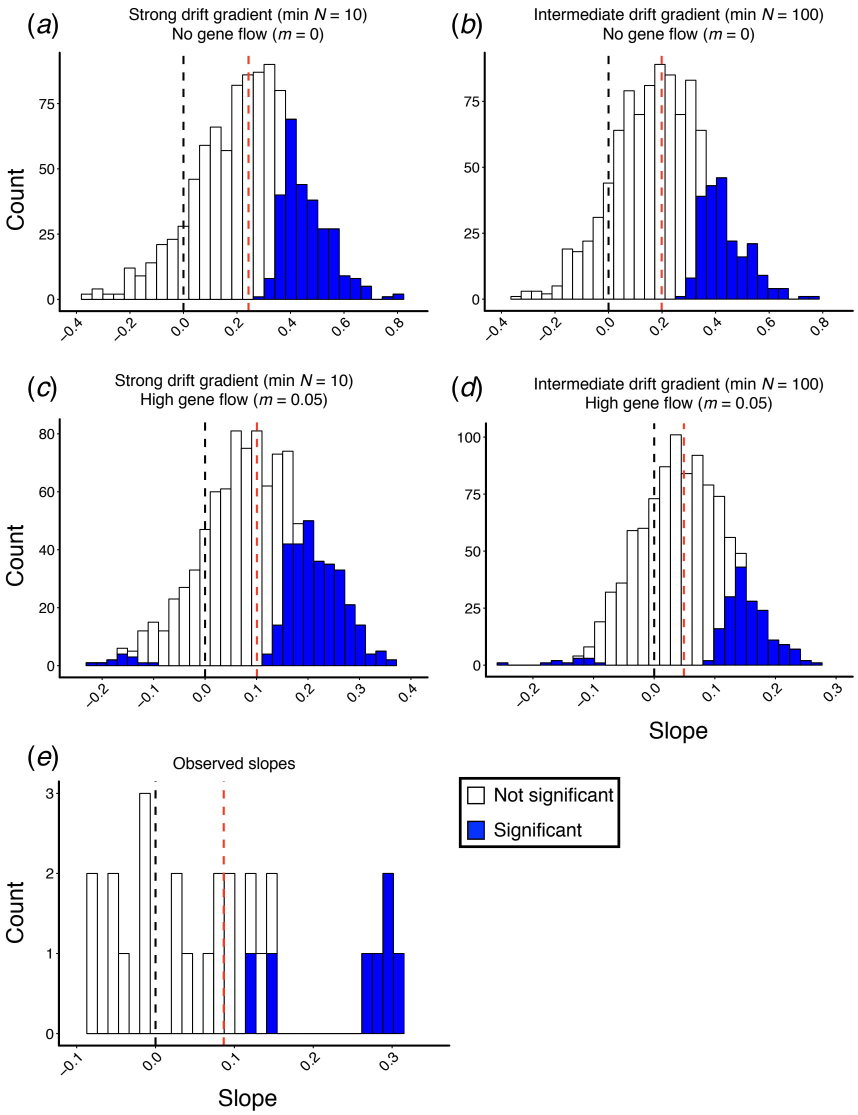


**Figure S13:** Distribution of standardized slopes for simulated (*a* through d) and observed (e) cyanogenesis clines. Simulated slopes were generated using drift scenario 1 (i.e. gradient in carrying capacity) under varying les of gene flow and strengths of drift. (*a*) Slopes from simulations under a strong gradient in drift (minimum urban $N = 10$) and no gene flow. (*b*) Slopes from simulations under an intermediate gradient in drift (minimum urban $N = 100$) and no gene flow. (*c*) Slopes from simulations under a strong gradient in drift (minimum urban $N = 10$) and high gene flow ($m = 0.05$). (*d*) Slopes from simulations under an intermediate gradient in drift (minimum urban $N = 100$) and high gene flow ($m = 0.05$). (*e*) Distribution of slopes from urban-rural cyanogenesis clines observed across cities (n = 26) by Thompson et al. (2016) and Johnson et al. (2018). Blue bars represent clines that are significant at $P < 0.05$. Black dashed bar over histograms represents a slope of zero whereas the red dashed bar represents the mean slope.

**Supplementary tables**

**Table S1:** Parameters used in simulations to manipulate the strength of drift, selection and gene flow among populations in the formation of spatial clines in HCN.

| Parameter | Description |
|---|---|
| Maximum amount of gene flow | Determines the maximum proportion of alleles (*CYP79D15* and *Li*) exchanged between any two populations. The actual proportion depends on the distance between populations (see text) |
| Maximum carrying capacity | Determines the carrying capacity of the largest habitat patch on the landscape (rural-most or urban-most population). |
| Minimum carrying capacity | Determines the carrying capacity of the smallest habitat patch on the landscape (rural-most or urban-most population). |
| Founder proportion | Proportion of alleles sampled when founding new populations. Lower proportions result in stronger effects of drift. This is equivalent to manipulating the number of individuals sampled to form new populations. |
| Maximum probability of creation of new population | Maximum probability that a new population is created. Actual probability depends on the population's size such that larger populations have a greater probability of creating new ones. Value is fixed at 1.0 so that populations at carrying capacity always found new populations. |
| Maximum selection coefficient | Maximum strength of selection acting on cyanogenic or acyanogenic genotypes. Actual strength of selection depends on a population's position in the landscape matrix. |
| Frequency of dominant *CYP79D15* | Initial frequency of the dominant allele at the *CYP79D15* locus. |
| Frequency of dominant *Li* | Initial frequency of the dominant allele at the *Li* locus. |
| Intrinsic rate of population increase | Intrinsic growth rate parameter used in logistic equation of population growth. Fixed at 1.5. |

**Table S2:** Details for all eight cases we simulated exploring the combined effects of drift, gene flow and selection on the formation of clines in HCN.

| Results figure | Mechanisms explored | Parameters controlling drift | Initial allele frequencies | Selection coefficients | Levels of gene flow |
|---|---|---|---|---|---|
| *Drift scenario 1: Gradient in maximum population size* | | | | | |
| 2*a, c* | Drift | Max $K$ = 1000 (Rural); Min $K$ = 10; 100; 500; 1000 (Urban) | *CYP79D15* = 0.5 *Li* = 0.5 | None | None |
| 2*b, d* | Drift, gene flow | Max $K$ = 1000 (Rural); Min $K$ = 10 (Urban) | *CYP79D15* = 0.5 *Li* = 0.5 | None | 0; 0.001; 0.0025; 0.005; 0.01; 0.02; 0.035; 0.05; 0.1; 0.2; 0.35; 0.5; 1.0 |
| S6 | Drift, gene flow, allele frequency variation‡ | Max $K$ = 1000 (Rural); Min $K$ = 10 (Urban) | *CYP79D15* = 0.1; 0.5; 0.9 *Li* = 0.1; 0.5; 0.9 | None | 0; 0.01; 0.05 |
| 3*c, d* | Drift, selection, gene flow† | Min $K$: 10 (Rural) Max $K$: 1000 (Urban) | *CYP79D15* = 0.5 *Li* = 0.5 | 0; 0.001; 0.0025; 0.005; 0.0075; 0.01; 0.025; 0.05; 0.1; 0.2 | 0; 0.01; 0.05 |
| *Drift scenario 2: Colonization through serial founder events* | | | | | |
| S3*a, b* and S4 | Drift, gene flow | **Founder proportion:** 0.01; 0.02; 0.035; 0.05; 0.075; 0.1; 0.2; 0.5; 0.75; 1.0 | *CYP79D15* = 0.5 *Li* = 0.5 | None | 0; 0.01; 0.05 |
| S7 | Drift, gene flow, allele frequency‡ | **Founder proportion:** 0.01; 0.2; 1.0 | *CYP79D15* = 0.1; 0.5; 0.9 *Li* = 0.1; 0.5; 0.9 | None | 0; 0.01; 0.05 |
| S5 | Drift, selection, gene flow† | **Founder proportion:** 0.01; 0.2; 1.0 | *CYP79D15* = 0.5 *Li* = 0.5 | 0; 0.001; 0.0025; 0.005; 0.0075; 0.01; 0.025; 0.05; 0.1; 0.2 | 0; 0.01; 0.05 |
| *No drift* | | | | | |
| 3*a, b* | Selection, gene flow | None. All populations with constant $K$ = 1000 | *CYP79D15* = 0.5 *Li* = 0.5 | 0; 0.001; 0.0025; 0.005; 0.0075; 0.01; 0.025; 0.05; 0.1; 0.2 | 0; 0.01; 0.05 |

† Modified drift scenarios. Refers to gradient in drift running from urban (weak drift) to rural (strong drift) rather than rural to urban, as simulated in other scenarios. These are used to explore drift-selection balance.
‡ Allele frequencies crossed factorially.

**Supplementary text**

Supplementary text S1 — Methods for drift scenario 2: Colonization and founder events

*Text S1a — Question 1: Does genetic drift influence the formation of spatial clines in HCN?*

The second scenario represents a case where the urban core is gradually colonized from surrounding rural regions. The simulations began with a single rural population at carrying capacity and adjacent patches were colonized toward the urban end until all patches contained populations (figure S8*b* and S8*c*). Given white clover's history of being seeded into agricultural pastures as forage and fertilizer [1], this represents a biologically realistic colonization scenario. Nonetheless, we emphasize that in the absence of selection, the direction of colonization is unimportant and was chosen to represent the direction of clines observed across urbanization gradients [2], and serves only to explore whether colonization through repeated founder events influences the formation of phenotypic clines in HCN. In later simulations (text S1b below), we reverse this colonization scenario to examine the formation of clines when drift and selection are in opposition.

There was no gradient in carrying capacity in this scenario; rather, the strength of drift was manipulated by varying the strength of founder events, determined as the proportion of alleles sampled from the parent population (i.e. smaller proportion = stronger founder event) and is equivalent to sampling a finite number of individuals. We initially simulated 10 different founding proportions (0.01; 0.02; 0.035; 0.05; 0.075; 0.1; 0.2; 0.5; 0.75; 1.0) to explore the formation of clines under a broad range of serial founder events. For each founding proportion, we simulated three levels of gene flow: $m$ = 0, 0.01, and 0.05, representing no, low, and high gene flow, respectively.

In our simulations, the probability that a population colonizes an adjacent patch depends on its size. The probability of colonization is 1.0 for populations at carrying capacity and decreases linearly with decreasing population size. Because founder events reduce the size of newly formed populations, serial founder events would result in populations becoming rapidly extinct (or exceedingly small), preventing the colonization of new patches. We therefore implemented a model of logistic population growth allowing populations to grow every generation until they reached carrying capacity. Under this model, a population of size 10 took 27 generations to reach a carrying capacity of 1000 (growth rate [r] = 1.5). The mean number of generations required for the landscape to become filled with populations ranged from 40 with no founder events (proportion = 1.0) to ~410 with strong founder events (proportion = 0.01, figure S2*a*). Similarly, following colonization, the mean number of generations required for the urban-most population to reach carrying capacity ($K$ = 1000) ranged from 0 with no founder events to ~30 with strong founder events (figure S2*b*). Simulations were run for 500 non-overlapping generations beyond when all patches on the landscape contained populations. These simulations thus explored the effects of drift—manipulated through serial founder events—at generating clines in HCN under varying levels of gene flow in the absence of selection.

*Text S1b — Question 3: What are the interactive effects of genetic drift and selection on the formation of clines in HCN?*

We sought to understand the combined effects of drift, gene flow and selection on the formation of clines in HCN, and specifically the extent to which selection can counter the formation of clines under drift. In this scenario, we manipulated drift in the same way described above (text S1*a*) but with colonization occurring from urban ($N$ = 1000) to rural ($N$ = 10) populations (rather than rural to urban as above). We simulated three founding proportions from among the 10 described above: 0.01, 0.2, and 1.0, representing strong, intermediate, and no effects of drift through founder effects, respectively. These were chosen as they sufficiently capture the variation in the effects of founding events on the formation of clines. Selection favoured HCN+ genotypes in rural populations and HCN– genotypes in urban populations, as described above. As such, the stochastic loss of dominant alleles in smaller rural populations is countered by their higher fitness. We additionally included three levels of gene flow: $m = 0$, 0.01, and 0.05, representing no, low, and high gene flow, respectively. These simulations enabled us to identify the strength of selection necessary to counteract the loss of HCN due to drift through serial founder events and examine the formation of HCN clines under opposing forces of drift and selection, with varying levels of gene flow.

<u>Supplementary text S2 — Results for drift scenario 2: Colonization and founder events</u>

*Text S2a — Question 1: Does genetic drift influence the formation of spatial clines in HCN?*

In the absence of selection. serial founder events during the colonization of urban populations overwhelmingly caused the formation of positive clines, although the results are more complex than those under a spatial gradient in carrying capacity (see results for "Question 1" in main text, figure 2). With no or low gene flow, the strength of clines reached a maximum of $\beta \approx 0.006$ with an intermediate founder size during colonization (founder proportion = 0.2, figure S3*a*), and declined as the founder size either increased or decreased from this point. However, high gene flow eliminated this effect, instead leading to a gradual decrease in mean cline strength from $\beta \approx 0.0016$ when drift was strong ($0.01 \leq$ founder proportion $\leq 0.035$) to $\beta \approx 0.0002$ when drift was weak (founder proportion = 1.0, figure S3*a*). Similarly, the proportion of significantly positive clines peaked at ~75% when drift was strong (founder proportion = 0.1), and decreased as the strength of drift increased or decreased from this point (figure S3*b*). By contrast, the frequency of significantly negative clines increased gradually from ~17% to ~25% as the strength of drift decreased (figure S3*b*).

The peak cline strength and proportion of significantly positive clines at intermediate founder effect strengths can be best understood by exploring the dynamics of HCN loss as the landscape is colonized. When founder effects were very strong (e.g. founder proportion = 0.01), HCN was lost so rapidly during colonization that reductions in HCN were non-linear (figure S4*a*), and clines were therefore only weakly positive ($\beta = 0.003$, dotted line in figure S4*b*). By contrast, when founder effects were absent (e.g. proportion of founding alleles = 1.0), HCN was never lost from the matrix (figure S4*a*) and clines were very weak as the frequency of HCN shows little change across space ($\beta = 0.0009$, dashed line figure S4*b*). However, when founder effects were of intermediate strength (e.g. proportion of founding alleles = 0.2), HCN was maintained for longer periods of time during colonization (figure S4*a*) and its frequency changes substantially across space, resulting in strongly positive, linear clines (solid line in figure S4*b*).

*Text S2b — Question 3: What are the interactive effects of genetic drift and selection on the formation of clines in HCN?*

Serial founder events from urban to rural populations constrained the ability of selection to generate strongly positive cyanogenesis clines. In the absence of founder events, increasing selection led to consistently stronger clines, independent of the level of gene flow (figure S5*a*). When gene flow was low or absent, intermediate founder events (founding proportion = 0.2) resulted in the mean strength of clines being negative for all $s < 0.025$ (figure S5*b*) whereas the strongest positive clines occurred when $s = 0.05$ ($\beta \approx 0.005$ for both low and no migration). High gene flow reduced the extent at which negative clines were formed by selection (figure S5*b*) and weaker selection ($s > 0.005$) was required before positive clines evolved in the presence of high gene flow. When founder effects were strong, selection had to be greater than 0.0025, 0.01, and 0.005 to generate positive clines when gene flow was absent ($m = 0$), low ($m = 0.01$) and high ($m = 0.05$), respectively (figure S5*c*). The strongest positive clines occurred when $s = 0.05$ ($\beta \approx 0.002$ for no and low gene flow). These results are consistent with intermediate founder effects generating the strongest clines in HCN and further demonstrate that strong selection is required to overcome the formation of clines in the presence of an opposing drift gradient.

Serial founder events also influenced the extent to which selection generated positive and negative cyanogenesis clines. In the absence of founder events, and when selection is less than 0.005, both positive and negative clines occur with approximately 30% frequency (figure S5*d*). However, when $s > 0.005$, the frequency of positive clines rapidly increases to 100%, whereas negative clines declines to 0% (figure S5*d*). In the presence of intermediate founder events, negative clines are consistently more frequent for all $s < 0.1$, consistent with founder events of intermediate strength preferentially generating clines in HCN. However, when $s \geq 0.1$, both positive and negative clines occur at less than 10% frequency (figure S5*e*). Finally, strong founder events result in little change in the frequency of positive clines, which fluctuate around 20% for all but the strongest selection coefficient ($s = 0.2$, figure S5*f*). By contrast, the frequency of negative clines rapidly decreases from 45% in the absence of selection to 0% when $s \geq 0.0025$, becoming less frequent than negative clines when $s \geq 0.005$ (figure S5*f*). These results further demonstrate that strong selection is required to overcome the formation of clines in HCN in the presence of opposing gradients in genetic drift.

Supplementary text S3: Effects of initial allele frequency variation on cyanogenesis cline formation

*Text S3a — Drift scenario 1: Gradient in carrying capacity across the matrix*

The initial frequency of both dominant alleles influenced the formation and strength of phenotypic clines in HCN. The strongest clines ($\beta = 0.006$) occurred when the frequency of both dominant alleles (i.e. *CYP79D15* and *Li*) was 0.5 (figure S6*a*). The weakest clines occurred when the frequency of one or both dominant alleles was low (i.e. 0.1; $0.0005 < \beta < 0.002$) whereas clines of intermediate strength occurred when either or both alleles were at high frequency (i.e. 0.9; $0.003 < \beta < 0.005$; figure S6*a*). These results hold regardless of the levels of gene flow; increasing gene flow reduced the strength of clines, regardless of initial allele frequencies (figure S6*a*).

The proportion of significantly positive clines was always greater than the proportion of negative clines, independent of initial allele frequencies. The frequency of significantly positive clines peaked at 30% when the frequency of both dominant alleles was 0.5, followed by cases when one or both alleles were at low frequency (i.e. 0.1; 11 < % < 16) and finally by cases where one or both alleles were at high frequency (i.e. 0.9; 16 < % < 22; figure S6*b*). Significantly negative clines were rare and only arose when the frequency of one or both dominant alleles was high (figure S6*b*).

*Text S3b — Drift scenario 2: Colonization and founder events*

The initial frequency of dominant alleles in the rural-most population influenced the strength and formation of clines through serial founder events. In the absence of founder events, initial allele frequencies had little effect on the strength of clines, which were on average very weak and near zero ($0.00002 < \beta < 0.0006$), independent of migration rate (figure S7*a*). When founder effects were of intermediate strength (founding proportion = 0.2), the strongest clines occurred when the initial frequency of one or both dominant alleles was 0.5 ($\beta = 0.005$ when $m = 0$) whereas the weakest clines occurred when one or both dominant alleles was at low frequency (e.g. 0.1, $\beta = 0.0005$ when $m = 0$, figure S7*b*). This result holds regardless of the level of gene flow (figure S7*b*). We found a similar pattern when founder effects were strong (founding proportion = 0.01), although clines were on average weaker: the strongest clines occurred when the frequency of both dominant alleles was at 0.5 and when gene flow was low ($m = 0.01$, $\beta = 0.002$, figure S7*c*). These results are consistent with those found under drift scenario 1 and demonstrate that, on average, the strongest clines occur when the initial frequency of both dominant alleles is 0.5.

The initial frequency of dominant alleles also influenced the extent to which serial founder events generated positive and negative clines. In the absence of founder effects (founder proportion = 1.0), the frequency of positive clines peaked at 43% when the initial frequency of both dominant alleles was 0.1 and declined gradually to a frequency of 29% when both alleles began at 0.9 (figure S7*d*). By contrast, when the initial frequency of initial alleles was low (i.e. 0.1), negative clines occurred at 18% frequency, increasing gradually to 29% when both alleles began at 0.9, thereby matching the proportion of positive clines. Under intermediate founder effects (founding proportion = 0.2), positive clines were consistently more common than negative clines, with the exception of when both dominant alleles were at high frequency (i.e. 0.9 figure S7*e*). When founder effects were strong (founding proportion = 0.01), positive clines were again more common than negative clines, but a substantial difference between the two only occurred when one dominant alleles was at high frequency or when both began at a frequency of 0.5 (figure S7*f*). Thus, independent of initial allele frequencies, positive clines tended to evolve more frequently through serial founder events during the colonization of urban environments.

Supplementary text S4: Modelling gene flow

Most simulations varied the amount of dispersal between populations across the landscape to explore the effects of gene flow on the formation of clines due to drift and selection (see table S2). We modelled gene flow according to a modified version of Wright's island model [3]. We began by defining the level of gene (e.g. $m$ = 0.05), which represented the maximum proportion of alleles exchanged between any two populations. However, the realized proportion of alleles exchanged between any two populations depends on the distance between them, such that closer populations exchange more alleles. Every generation, each population could exchange alleles with all others in the landscape and thus the proportion of alleles immigrating into any one population depended on the mean proportion of immigrating alleles, averaged across all existing populations. Specifically, the frequency of the dominant allele (i.e. *CYP79D15* or *Li*) in population $x$ in the next generation ($p_{A_x,t+1}$) is given as:

$$p_{A_x,t+1} = p_{A_x,t}\left(1 - \overline{m_{w_x}}\right) + (\overline{p_{A_{wx},t}})(\overline{m_{w_x}})$$

where $p_{A_x,t}$ is the frequency of the dominant allele at *CYP79D15* or *Li* in population $x$ in the current generation, $\overline{m_{w_x}}$ is the weighted-mean immigration rate from all populations into population $x$ and $\overline{p_{A_{wx},t}}$ is the weighted-mean frequency of the dominant allele in the current generation for population $x$'s migrant pool, averaged across all other existing populations, respectively. Gene flow is assumed to decline linearly with increasing distance between populations such that there is effectively no gene flow between populations 1 and 40 of the landscape. Levels of gene flow and dominant allele frequencies were weighted by population size such that larger populations contributed more alleles to the migrant pool. Specifically, the weighted-mean immigration rate from all populations into population $x$ was calculated as:

$$\overline{m_{w_x}} = \frac{\sum_{y=1}^{n} m_{xy} \times N_y}{\sum_{y=1}^{n} N_y}$$

where $m_{xy}$ is the realized rate of gene flow between populations $x$ and $y$, based on the distance between them, $N_y$ is the size of population $y$, and $n$ represents the number of populations minus one (i.e. 39) since populations do not exchange migrants with themselves. Similarly, the weighted-mean dominant allele frequency for population $x$'s migrant pool was calculated as:

$$\overline{p_{A_{wx},t}} = \frac{\sum_{y=1}^{n} p_{A_y} \times N_y}{\sum_{y=1}^{n} N_y}$$

where $p_{A_y}$ is the frequency of the dominant allele in population $y$. We performed the above process separately for both dominant alleles (i.e. *CYP79D15* and *Li*) every generation.

Supplementary text S5: Effects of selection on linkage between *CYP79D15* and *Li*

We performed a small-scale simulation to examine the build-up of linkage disequilibrium (LD) between *CYP79D15* and *Li* when selection is acting on HCN. We initialized a single population with the frequency of both dominant alleles set to 0.5. From these allele frequencies, we calculated the frequency of all 16 possible diploid genotypes, assuming Hardy-Weinberg equilibrium. We then subjected these genotypes to selection, which acted against (negative selection coefficients) or in favor (positive selection coefficients) of cyanogenic genotypes. From the selected genotypes, we calculated the frequency of gametes, where heterozygotes were assumed to produce equal frequencies (i.e. 0.25) of all 4 possible gametes given the absence of physical linkage between *CYP79D15* and *Li* (i.e. recombination = 0.5). We calculated linkage disequilibrium as:

$$r^2 = \frac{D^2}{p_{CYP79D15} \times q_{CYP79D15} \times p_{\mathrm{Li}} \times q_{\mathrm{Li}}}$$

In this equation, $p_{CYP79D15}$, $q_{CYP79D15}$, $p_{\mathrm{Li}}$, and $q_{\mathrm{Li}}$ represent the frequency of dominant (i.e. p) and recessive (i.e. q) alleles at *CYP79D15* and *Li*. D represents the coefficient of linkage disequilibrium and is a function of gamete frequencies in any one generation and is calculated as:

$$D = (x_{11})(x_{22}) - (x_{12})(x_{21})$$

where $x_{11}$ represents the frequency of gametes composed of homozygous dominant alleles at *CYP79D15* and *Li*, $x_{22}$ the frequency of gametes composed of homozygous recessive alleles at *CYP79D15* and *Li* and $x_{12}$ and $x_{21}$ the frequency of gametes composed of heterozygous alleles at one or the other locus. Thus, $r^2$ is a measure of LD that accounts for allele frequencies and has a value of 1 when loci are in complete LD and 0 when they are in linkage equilibrium (i.e. independent of one another, [4] pp. 81-85). Gamete frequencies in following generations were then calculated from selected genotypes and this process was repeated recursively for 500 generations, allowing us to track the build-up of LD due to selection for cyanogenic and acyanogenic genotypes.

Our results show that the build-up of LD is minimal and decays rapidly over 500 generations (figure S10). Even under strong selection (e.g. -0.1), $r^2$ reaches a maximum just under 0.0005, which is sufficiently close to zero to consider the loci in linkage equilibrium.

Given these results, we ignored the build-up of LD due to selection in our simulations.

Supplementary text S6: Effects of generation chosen for analysis on the formation and strength of clines in HCN

The generation chosen for analyses had little effect on our ability to assess the contributions of drift, gene flow, or selection on the formation and strength of phenotypic clines in HCN. For simplicity, we demonstrate this only for a strong gradient in carrying capacity (drift scenario 1, maximum rural $N = 1000$, minimum urban $N = 10$) under varying levels of gene flow. Following the formation of a cline, differences in the mean strength of clines across varying levels of gene flow remain consistent, independent of generation (figure S11*a*). The only exception to this is the strength of clines in the absence of gene flow, which decreases gradually over time due to drift, resulting in weaker clines at generation 500 than is evident at generation 250 (figure S11*a*). Nonetheless, this has no effects on our interpretation that increasing the amount of gene flow reduces the mean strength of clines (see main text figure 2*b*). Similarly, differences in the proportion of significantly positive clines remain qualitatively similar across generations (figure S11*b*). Therefore, the generation chosen for analysis has no influence on our ability to interpret the role of migration in influencing the formation and strength of cyanogenesis clines formed via drift.

Supplementary text S7: Comparison of simulated slopes to standardized slopes for cyanogenesis clines observed across urban-rural gradients

We were interested in comparing the strength of clines produced by drift in our simulations to the strength of clines observed across urban-rural gradients in natural populations. For simplicity, we only examined the strength of clines simulated under weak or intermediate gradients in carrying capacity (drift scenario 1) and for no and high gene flow. Data for observed clines were obtained from [2] and [5]. Because the length of transects in our simulations and across cities varied, it was necessary to first standardize slopes before comparison. We standardized transects to a minimum value of 0 (urban-most population) and a maximum value of 1 (rural-most population). We then performed a linear regression using within-population HCN frequency as the response variable and standardized distance value as the predictor variable. Positive slopes represent less HCN in urban populations whereas negative slopes represent the opposite.

We found that the strength of observed clines is consistent with the strength of clines generated by drift in our simulations. The strongest simulated clines occurred under a strong gradient in drift in the absence of gene flow ($-0.35 < \beta_{simulated} < 0.81$, figure S13*a*). Increasing the amount of gene flow or decreasing the strength of the gradient in drift reduced the maximum strength of clines (figure S13*b* through S13*d*). The weakest simulated clines in HCN occurred under an intermediate gradient in drift with high gene flow ($-0.24 < \beta_{simulated} < 0.27$, figure S13*d*). The strength of observed clines ranged from –0.08 to 0.3 (figure S13*e*); thus, observed clines are within the range of even the weakest clines simulated under a gradient in drift, suggesting that drift is sufficient to generate clines as strong as those observed across replicated urbanization gradients.
